# Percolation transition determines protein size limit for passive transport through the nuclear pore complex

**DOI:** 10.1101/2021.12.17.473237

**Authors:** David Winogradoff, Han-Yi Chou, Christopher Maffeo, Aleksei Aksimentiev

**Affiliations:** Department of Physics, University of Illinois at Urbana-Champaign, Urbana, IL 61801, USA; Center for the Physics of Living Cells, University of Illinois at Urbana-Champaign, Urbana, IL 61801, USA; Beckman Institute for Advanced Science and Technology, University of Illinois at Urbana-Champaign, Urbana, IL 61801, USA

## Abstract

Nuclear pore complexes (NPCs) control biomolecular transport in and out of the nucleus. Disordered nucleoporins in the complex”s central pore form a permeation barrier, preventing unassisted transport of large biomolecules. Here, we combine coarse-grained simulations of an experimentally-derived NPC structure with a theoretical model to determine the microscopic mechanism of passive transport. Brute-force simulations of protein diffusion through the NPC reveal telegraph-like behavior, where prolonged diffusion on one side of the NPC is interrupted by rapid crossings to the other. We rationalize this behavior using a theoretical model that reproduces the energetics and kinetics of permeation solely from statistical analysis of transient voids within the disordered mesh. As the protein size increases, the mesh transforms from a soft to a hard barrier, enabling orders-of-magnitude reduction in permeation rate for proteins beyond the percolation size threshold. Our model enables exploration of alternative NPC architectures and sets the stage for uncovering molecular mechanisms of facilitated nuclear transport.

---

The nucleus secludes the genetic material of a eukaryotic cell to ensure the fidelity of transcription, replication and gene regulation processes. ^1–5^ Nuclear pore complexes (NPCs) perforate the nuclear envelope, a double lipid bilayer that encases the nucleus, providing a passage for biomolecular traffic in and out of the nucleus. Water, ions and small biomolecules, up to ∼5 nm in diameter, can pass through an NPC largely unimpeded. Unassisted transport of larger molecules is blocked,^6–8^ but, when combined with nuclear transport factors, cargoes up to 40 nm in diameter can traverse an NPC,^9,10^ fueled by a RanGTP/GDP cycling system. ^11,12^ The selectivity of nuclear pore transport is attributed to the properties of nucleoporin proteins (nups) that form a barrier to diffusion through the NPC”s central channel.^13,14^ Many of such nups have repeating phenylalanine-glycine (FG) motifs^15^ that form hydrophobic, highly dynamic,^16^ and intrinsically disordered domains^17^ known to interact with nuclear transport factors.^18,19^ Medically, disturbances in nuclear transport can lead to a number of human diseases, including cancer, viral infections, and neurodegenerative conditions. ^20,21^ Nuclear transport is also a key target for emerging gene therapy.^22^

Essential to the regulatory role of the NPC, its central mesh serves as a filter cutting off passive transport through the NPC above some molecular weight or geometrical size threshold. Theoretical studies have investigated passive diffusion through model narrow channels, showing that the presence of an attractive site within the channel can greatly increase the diffusive flux^23^ and enable transport selectivity. ^24^ For a given concentration of solutes, the attractive interactions can be tuned to optimize the transport,^25^ although, at high concentrations, the highest flux is achieved when an attractive potential is replaced by a barrier.^26^ The transport becomes even more nuanced when multiple binding sites are present within the channel.^27^ Relating to nuclear transport specifically, early experiments ^28,29^ observed proteins as large as 10 nm in diameter to passive diffuse across an NPC. More recent experiments^6,8,30,31^ established an absence of a sharp size threshold for the passive diffusion, although 5 nm diameter^30^ or a 40 to 60 kDa mass are often cited as effective size and molecular weight thresholds, respectively. Fluorescent tracking experiments revealed that, given enough time, proteins as large as 200 kDa and 8 nm in diameter can cross an NPC unassisted by transport factor,^6^ implying that passive transport of large cargoes through an NPC is not impossible, but is rather impractical, from a cell biology point of view.

Computational studies play an important role in evaluating possible mechanisms of nuclear pore transport by providing insight into microscopic processes not readily accessible to experiment. Molecular transport through the NPC has been examined within the framework of the kinetic theory,^32^ showing how selectivity can arise from the competition for the limited space inside the channel. Polymer theory methods have been applied to evaluate the effect of the cargo size on the configurational entropy of a polymer brush grafted to a model channel,^33^ suggesting that the size-dependent selectivity may originate from an entropic effect. Free-energy models have been developed to show how molecularly divergent NPCs in different biological species can perform essentially the same function, ^34^ how a complex free-energy profile may arise from electrostatic and hydrophobic residues within the NPC mesh, ^35^ including local electrostatic polarization of the nup domains. ^36^ Coarse-grained (CG) molecular dynamic simulations have been combined with a theoretical model to determine the effect of cohesive interactions on protein transport,^37^ characterize passive transport through a model NPC system, ^8^ and to directly evaluate the free-energy barrier for translocation through a synthetic channel decorated with disordered nup proteins,^38,39^ finding the local interaction with the mesh to enhance transport of larger particles.^40,41^

Here, we combine a computational model of an experimentally-derived NPC structure with a percolation transition analysis to determine the physical origin of the barrier to passive diffusion of globular proteins across an NPC. Through brute-force simulations of passive diffusion, we directly characterize the effect of protein size on the passive diffusion rate. We rationalize the observations by examining an ensemble of configurations realized by the nup mesh in the absence of any proteins, arriving with a general method for estimating the free-energy barrier and the transport rates. Using our theoretical framework, we discover a crossover in the scaling behavior of the passive diffusion rate on the protein size and identify percolation transition to be at the origin of the crossover. Our work sets the stage for computational characterization of passive transport through NPC variants differing in their composition, stoichiometry and physical dimensions, providing a means to connect variations in NPC structure^42,43^ and plasticity^44,45^ to the NPC”s function as the gate keeper of nuclear transport.

## Results and discussion

### Computational model of experimentally-derived NPC structure

Starting from a composite structure of a human NPC, ^46^ we developed a computational model that included a custom grid-based potential representation of the nuclear envelope and of the protein scaffold and a one-amino-acid-per-bead representation of the disordered FG-nup mesh, Fig. 1a,b. The shape of the nuclear envelope was derived from a cryo-electron tomography reconstruction (EMD-3103)^47^ whereas the scaffold potential was derived from the composite structure. ^46^ Matching the stoichiometry of a composite NPC structure, ^46^ our computational model included 32 copies of each Nsp1, Nup49, Nup57, Nic96, and Nup145N proteins, with their disordered mesh parts generated as a self-avoiding random walk starting from the point anchoring each nup to the scaffold. We chose not to include cytoplasmic filaments or the nuclear basket in our model, as their structures remains poorly characterized by experiment. The model was simulated using an in-house developed software ARBD. ^48^ The interactions between the beads representing the disordered mesh was described using the model developed by the Onck laboratory.^49,50^ Custom potentials described interactions of the FG-nup beads with the NPC scaffold and the nuclear envelope, see Methods for details. Two 7.5 millisecond replica simulations of our computational model characterized the highly heterogenous and dynamic ensemble of conformations adopted by the network of disordered FG-nup proteins. Figure 1a and Supplementary Movie 1 illustrate one simulation trajectory. The simulations revealed formation of transient channels within the mesh network, connecting the two compartments on the opposite sides of the NPC complex. One such channel is clearly visible in the 5,000 *µ*s overhead snapshot in Fig. 1a. The local density of the FG-nup mesh, averaged over the two simulation trajectories and over the symmetry of the NPC complex, Fig. 1b, displays a donut-shaped region of high (*>*55 mg/mL) local density surrounding the scaffold”s inner ring, and reduced density within the central channel. Averaging the local densities of individual nups, Fig. 1c, we find Nsp1, Nup49, Nic96 and Nup57 to extend into the central channel, with Nsp1 extending the farthest. At the same time, Nup145N is seen to primarily fill the space between the inner and outer rings of the NPC, whereas Nic96 partially fills the cavities present in the structured protein scaffolding, consistent with its known role in holding the NPC assembly together.^51^

**Fig. 1:**
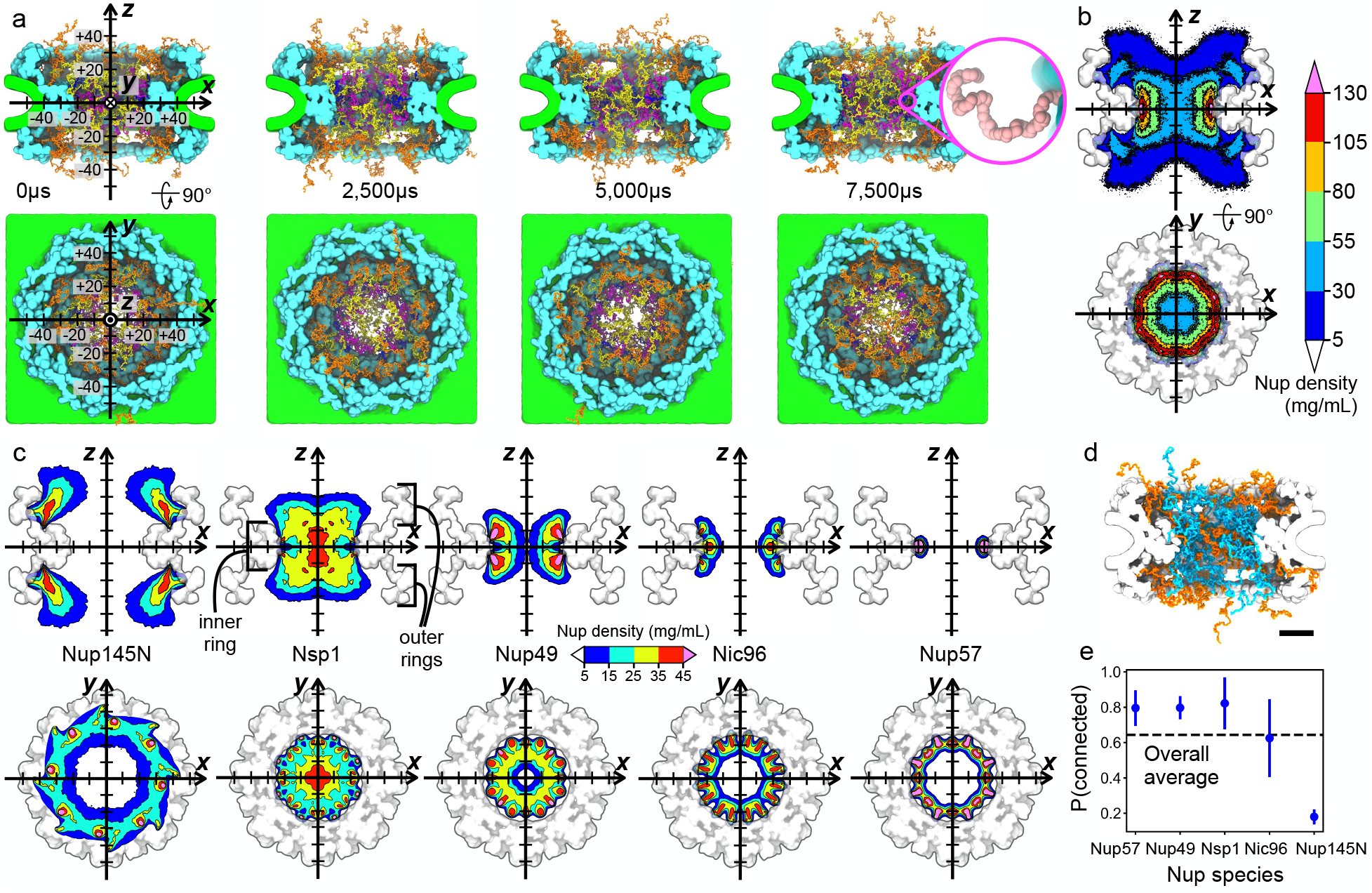
Equilibrium structure of a complete NPC complex. **a** Equilibration simulation of the NPC model. The computational model consist of a nuclear envelope (green), structured protein scaffold (cyan), and disordered FG-nups mesh consisting of 32 copies of each Nup145N (orange), Nsp1 (yellow), Nup49 (magenta), Nic96 (dark blue) and Nup57 (pink). Labels for the coordinate system (on the left) are in nm. Inset shows a magnified view of Nup57; each bead represents one amino acid. **b** Cross sections of the average FG-nup amino acid density. The 3D density map was generated by averaging instantaneous configurations of the computational model every 0.1 *µ*s over the final 6-ms fragments of two independent 7.5-ms simulations. The side and top view cross sections were additionally averaged along the *y* and *z* coordinate, respectively, within the [− 7.5, +7.5] nm range. **c** Average density maps of individual FG-nup species generated using the same protocols as the density map shown in panel b. Black brackets define the “inner ring” and “outer rings” regions of the NPC scaffold. **d** Representative configuration of the NPC mesh, where individual FG-nups making at least one contact with another FG-nup are shown in blue and those without such contacts are shown in orange. Scale bar, 20 nm. **e** The fraction of nups forming at least one interchain contact, by species. Error bars represent standard deviation. The dashed line shows the fraction averaged over all species. An interchain contact was defined as having two residues from different chains within 0.8 nm of one another.

A property of the FG-nup mesh that is often discussed with regard to the mechanism of nuclear transport is the extent to which nups connect together to generate a mesh. Analysis of our simulation trajectories shown that about 65% of all FG-nups chains are connected, on average, forming at least one contact with a residue from another nup. A representative configuration of the FG-nup mesh, Fig. 1d, shows that the nups tethered to the NPC”s inner ring are almost all connected: over 80% of Nup57, Nup49 and Nsp1 form at least one contact with another nup, Fig. 1e. Nup145N is the least likely to form interchain contacts with other nups because of its anchor position that is far from the midplane dividing the NPC in two halves along its pore axis, whereas Nic96 exhibits the greatest variability of its connectedness. In all, a single nup can form up to twenty interchain contacts, Supplementary Fig. 1a. The differences among nup species is consistent with Nup145N and Nic96”s classification as “adaptor nups,” distinct from Nsp1, Nup49 and Nup57, which are considered to be the “channel nups.” Hydrophobicity is hypothesized to play an important role in forming the FG-nup mesh. Interestingly, most of the interchain contacts observed in our simulations were formed between a hydrophobic and a polar residue (∼45%), about twice as likely as a hydrophobic–hydrophobic or polar–polar contact, which each occurred about 22% of the time, see Supplementary Fig. 1b. Our results suggest that FG-nups within the NPC central channel are mostly connected with each other, but these interactions are transient and are not necessarily prescribed by the local hydrophobicity of the nup residues.

### Passive diffusion of individual proteins across the NPC

To examine the process of passive diffusion across the NPC, we constructed 26 simulation systems each containing, in addition to our computational model of the NPC, a single protein represented as a rigid body that interacted with the NPC via the same potentials as beads of the FG-nup mesh, see Methods for details. Thirteen unique protein species were modeled, ranging in molecular mass from 5 to 145 kDa, Fig. 2a and Supplementary Table 1. To increase the probability of a protein”s encounter with the FG-nup mesh, the center of mass (CoM) of the protein was subject to a confinement potential, Fig. 2b, which reduced the volume available for protein diffusion to a cylinder co-axial with the pore of the NPC. Our rigid body approximation of the protein preserved the protein shape at one-bead-per-residue resolution, Fig. 2c. The translational and rotational diffusion constants of each protein were separately defined along the principal component axes of the corresponding rigid-body and used to simulate translational and rotational displacements via a Brownian dynamics algorithm. Two independent 6,000 *µ*s simulations were performed for each protein species, differing only by the initial location of the diffusing protein, Fig. 2b. The protein diffusion simulations were also repeated using a narrower, 25-nm-radius confinement potential, Supplementary Fig. 2.

**Fig. 2:**
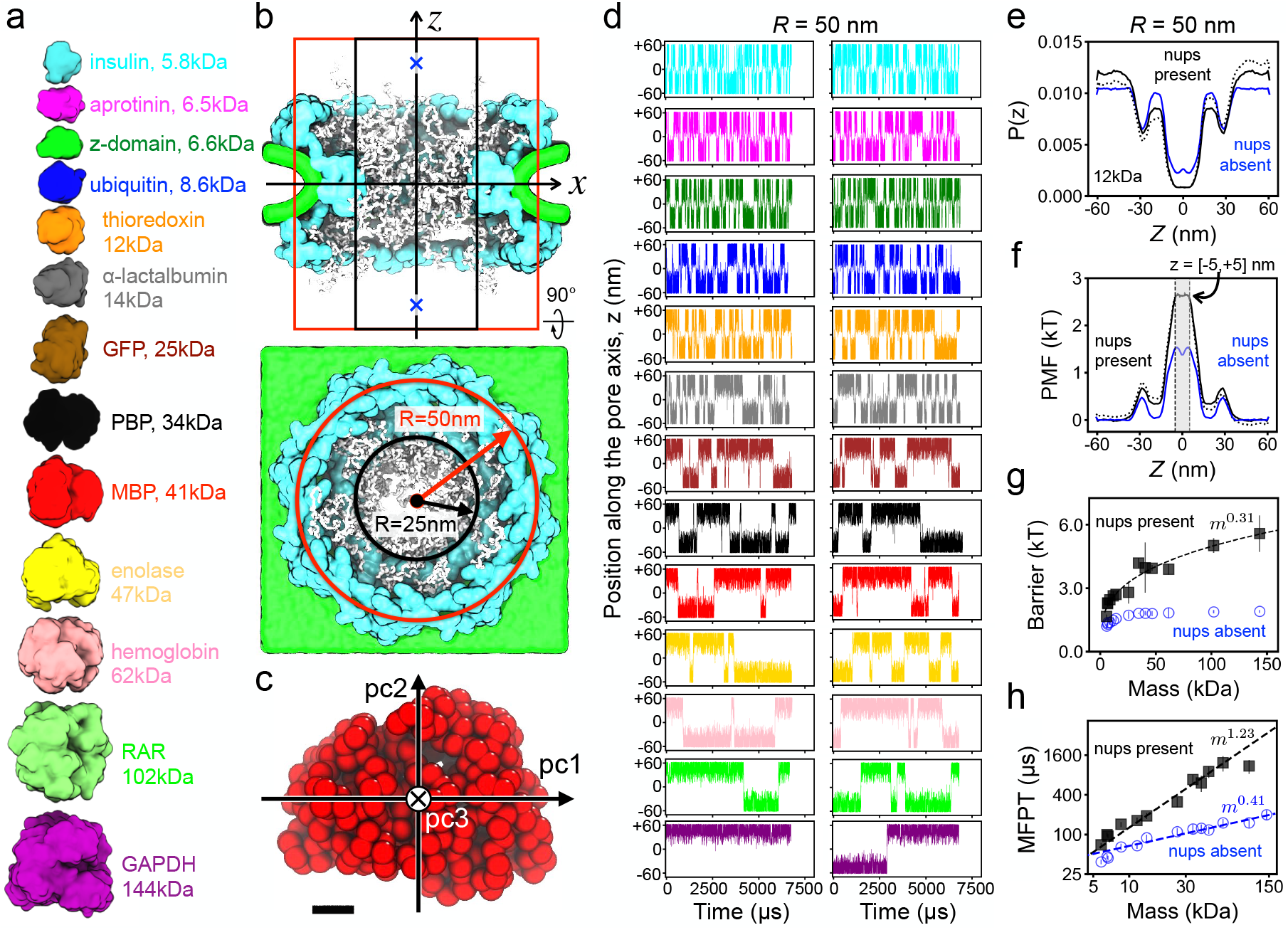
Passive diffusion across the NPC. **a** To-scale structural representation of all proteins used for CG simulations of passive diffusion. **b** Side (top) and overhead (bottom) view of the simulation systems. Black and red lines indicate the approximate location of a cylindrical confinement potential of 25 and 50 nm radius, respectively. Blue crosses mark the two initial locations of the protein (at *z* = ± 50 nm). **c** Rigid-body model of a maltose binding protein (MBP) where each bead represents one amino acid. Pc1, pc2 and pc3 denote the principal axes of the protein. Black scale bar, 1 nm. **d** Center-of-mass *z* coordinate of each protein (colors defined in panel a) versus simulation time. The simulations traces in the two columns differ by the initial placement of the protein. The simulation were performed in the presence of a 50 nm-radius confinement potential. **e** Normalized distribution of the CoM *z* coordinate of thioredoxin. The black dotted and solid lines show the distribution extracted directly from the simulations and the symmetrized distribution, respectively. The blue line shows a symmetrized distribution for the simulation carried out in the absence of the FG-nup mesh. **f** Potential of mean force (PMF) for thioredoxin transport across the NPC. A PMF barrier is defined as the average value within |*z* |*<* 5 nm. **g** PMF barrier versus protein molecular mass determined from CG simulations of protein diffusion through our complete NPC model (black squares) and the model missing all FG-nups (blue circles). Error bars are defined as the average point-by-point difference of the unsymmetrized PMF values from − 50 *< z <* 0 nm and 0 *< z <* 50 nm intervals. Line shows a power law fit to the data. **h** Mean first-passage time (MFPT) versus protein molecular mass, both axes logarithmic. Power-law fits are shown as dashed lines in panels g & h. In panel h only, the power-law fit to the “nups present” data included proteins up to 62 kDa (hemoglobin).

In a typical simulation, a protein was observed to translocate from one side of the NPC to the other multiple times as the FG-nup mesh continuously changed its conformation, see Supplementary Movie 2. The plots of the proteins” CoM *z* coordinate versus simulation time, Fig. 2d, reveal a telegraph-like behavior, where prolong intervals of protein diffusion within one of the compartments are interrupted by rapid translocations through the FG-nup mesh to the other compartment. The rate of spontaneous transport from one compartment to the other visibly decreases as the protein mass increases, the proteins are also less likely to approach the NPC”s midplane (*z* = 0 nm). For each successful translocation event, there are many more half-way spikes, indicating unsuccessful translocation attempts. Over the aggregate simulation time of 12,000 *µ*s, the largest protein, GAPDH, underwent only one successful translocation and, hence, simulations of larger proteins were not conducted.

We rationalize the protein translocation traces by computing, for each protein species, the distribution of the protein”s CoM *z* coordinate, *P* (*z*). Assuming our simulations have adequately sampled the configurational space, we can interpret a Boltzmann inversion of the *z* coordinate distribution, −*k*_B_*T* ln *P* (*z*), as a potential of mean force (PMF) acting on the protein as it passes through the NPC. Figure 2e,f shows a representative normalized distribution and the PMF, respectively, for one protein; Supplementary Fig. 4 and Supplementary Fig. 5 show similar data for all protein species. For comparison, protein diffusion simulations were performed using an NPC model that lacked the disordered FG-nup mesh. According to the *P* (*z*) and PMF plots, the primary barrier to passive diffusion lies at the NPC”s midplane, the location of NPC scaffold”s inner ring. The two smaller secondary barriers correspond to the scaffold”s outer rings. Accordingly, the secondary peaks are absent in the PMFs extracted from the simulations carried out under a narrower confinement potential, Supplementary Figs. 6 & 7.

Having defined the PMF barrier as the average value of the PMF profile within 5 nm of the NPC midplane, Fig. 2f, we investigate how the barrier height depends on the protein mass, Fig. 2g. For small proteins (*<*10kDa), the protein scaffold is seen to contribute a considerable fraction of the translocation barrier, which can be appreciated from a comparison of the PMFs obtained with and without the FG-nup mesh, Fig. 2f. The effect of the mesh becomes more dominant for larger proteins and the translocation barrier increases with the protein mass, *m*, as ∼ *m*^0.31^. In the case of protein transport through nup-less NPC, the translocation barrier varies considerably less with the protein mass, as ∼ *m*^0.11^, reflecting a modest change in the available configurational space because of the steric interactions with the NPC scaffold. Conversely, noticeably smaller barriers were extracted from the simulations carried out under a narrower, 25 nm-radius confinement potential, Supplementary Fig. 2, as the scaffold occupied a smaller fraction of the simulation volume.

We characterize the protein translocation time by recording the time elapsed from the moment the protein first enters the NPC volume (defined to be at *z* = ±20 nm) to the moment the protein exits the NPC volume on the opposite side, i.e., the first-passage time. Multiple translocations were observed for each protein during our CG simulations (except for GAPDH) and averaging over all translocation events gave the mean first-passage time (MFPT). In the absence of nups, the MFPT is observed to increase with the protein mass as *m*^0.41^, Fig. 2h, or as *m*^0.44^, Supplementary Fig. 2f, depending on the width of the confinement potential. This scaling exponent is close to but slightly higher than the free diffusion limit (*m*^1*/*3^), which we attribute to small yet not negligible effect of the NPC scaffold. A much steeper dependence was observed for a complete NPC model, with the MFPT increasing as *m*^1.23^, Fig. 2h, or as *m*^0.97^, Supplementary Fig. 2f. Thus, the transport rate in the absence of FG-nups appears to scale, approximately, with the protein radius (∼ *m*^1*/*3^), and, in the presence of nups, with the protein volume (∼ *m*^1^) for the range of masses explored by our CG simulations.

An experimental study that measured the accumulation of dye-labeled GFP dimer (about 50 kDa) into the nucleus of permeabilized HeLa cells^52^ provides us the opportunity to compare our simulated transport rates to experiment. Under a 50 nm-radius confinement, the protein concentration in our simulations is about 5 *µ*M. Linearly extrapolating from the experimentally measured transport rate to a GFP dimer concentration of 5 *µ*M, the expected experimental transport rate is ∼300 molecules/s for one NPC, which corresponds to a mean first-passage time of about 3.3 ms. In our CG simulations, mean first-passage time of a protein of a comparable size (enolase, 47 kDa) is about 0.9 ms Fig. 2h, which is a factor of four faster than expected from experiment. Thus, the simulated timescale of the NPC transport is of the same order as in experiment.

### Free diffusion determines the timescale of successful crossings

The telegraph-like shape of the protein permeation traces, Fig. 2d and Supplementary Fig. 2b, suggests that the timescale of a successful crossing event is orders of magnitude smaller than the timescale separating successful crossings. The latter is expected to depend on the effective concentration of the protein and, indeed, we observe about a three-fold decrease in the number of crossing events when changing the radius of the confinement potential from 25 to 50 nm, slightly less than the expected factor of four because a part of the additional volume of the larger confinement potential is occupied by the NPC”s scaffold. Figure 3a and Supplementary Movie 3 illustrate one representative crossing of a maltosebinding protein (MBP), which completes in about 6 *µ*. The passage of the MBP along the pore axis is accompanied by a similar magnitude displacement perpendicular to the pore axis. The configuration of the FG-nup mesh rearranges significantly over the course of the crossing event. The MBP permeation trace, Fig. 3b and c, shows that the MBP”s position during the crossing, i.e., from 5331 to 5337 *µ*s, decreases almost monotonically in *z*, after a partial (failed) crossing immediately beforehand. A zoomed-in view on another crossing event, Fig. 3d, from the same simulation show significant back-tracking but, ultimately, the protein crosses the NPC in less time than in the event shown in Fig. 3c. A careful look at the simulation trace, Fig. 3b, reveals that when a protein enters the NPC volume (defined as |*z*| *<* 20 nm), the protein almost always fails to complete the crossing, although the relative rate of success and failure depends on the protein size.

**Fig. 3:**
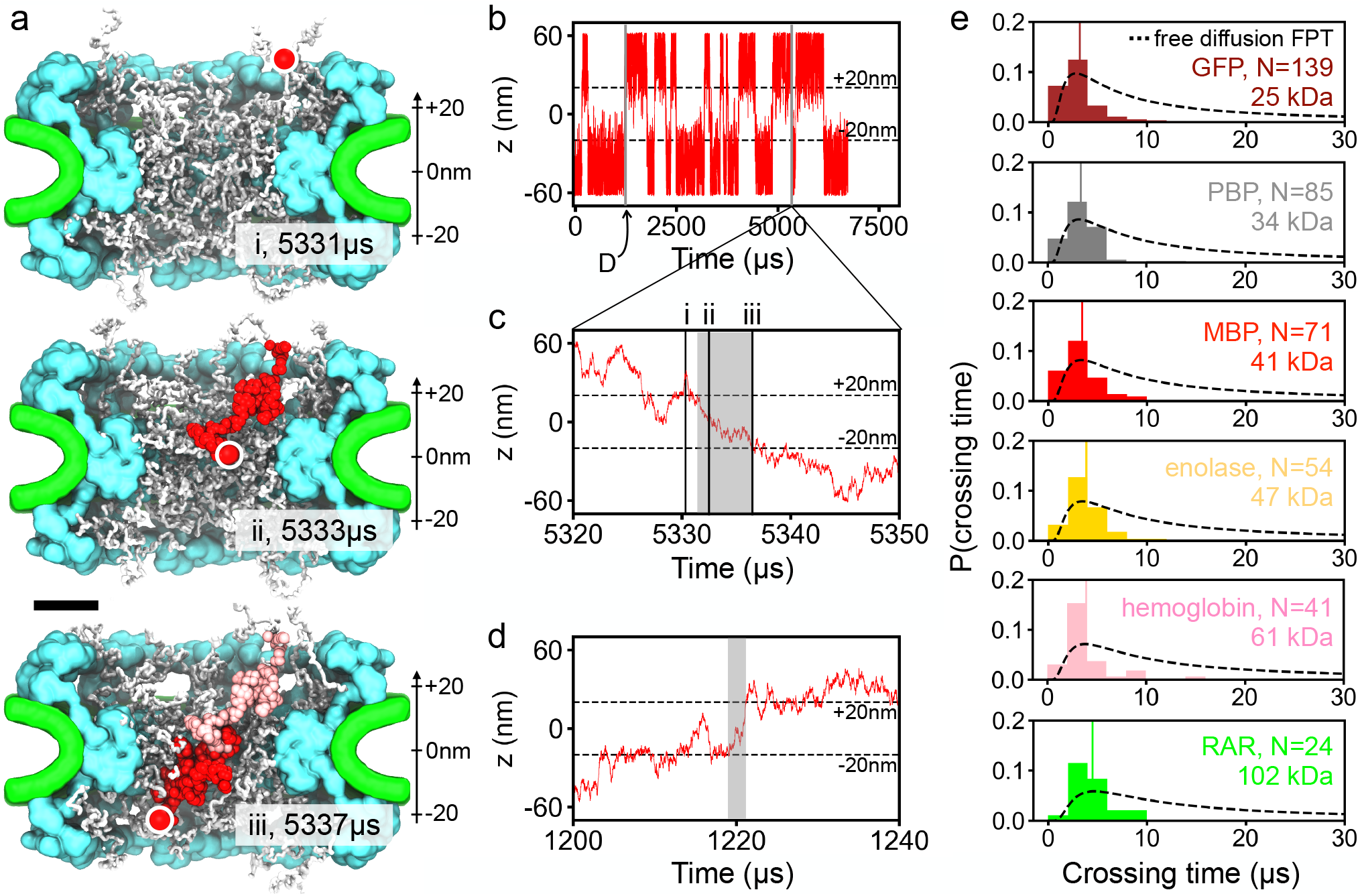
Timescale of protein crossings. **a** Successful translocation of a maltose-binding protein (MBP, red) through the NPC. Three instantaneous configurations of the NPC are shown using white for the FG-nups, cyan for the protein scaffold and green for the lipid bilayer. The circle indicates the location of the MBP protein at each configuration. Red and pink beads illustrates the configurations explored by the protein between the instantaneous configurations (sampled every 25 ns). Black scale bar, 20 nm. **b** CoM *z* coordinate of MBP simulated under a 25 nm radius confinement potential. Two crossing events form this trace are shown in detail in panels c and d. **c** Zoomed-in on the crossing event trace. The same event is illustrated by snapshots (i, ii, iii) in panel a. **d** Example of another crossing event. The gray rectangles in panels c and d illustrate the time interval defined as a crossing time for the analysis shown in panel e. **e** Distribution of crossing times for six protein species. *N* specifies the total number of crossing events used to construct each histogram. Each normalized histogram was constructed using 15 evenly-spaced bins, from 0 to 30 *µ*s, and the crossing time data from both confinement potential simulations, the average of each shown as a vertical line. The dashed lines show the distributions of first-passage time for a freely diffusing particle of the same diffusion constant as that of the corresponding protein.

To characterize the timescale of the crossing events, we define a crossing time as the time elapsed from the moment the protein exists the NPC volume (at *z* = ±20 nm) to the last prior moment the proteins crossed the NPC boundary on the other side of the NPC, see Fig. 3c,d for two examples. Note that this definition differs from that of a first passage, as the crossing time does not account for the time the protein spends meandering on the entrance side of the NPC volume after crossing its boundary for the first time. Using the above definition of the crossing time, we collected crossing time statistics for six proteins ranging in their molecular mass from 25 to 102 kDa. The average crossing time is seen to shift towards the right, though its molecular mass dependence is considerably less pronounced that that of the MFPT, Fig. 2h and Supplementary Fig. 2f, and appears to roughly follow the dependence of the diffusion constant on the protein mass (average crossing time ∼ *m*^0.26^ whereas 1*/D* ∼ *m*^1*/*3^). For reference, we plot in Fig. 3e theoretical distributions,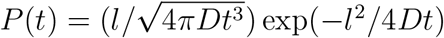, of the time, *t*, required for a particle of the same diffusion constant as that of the corresponding protein to travel distance *l* = 40 nm along a straight line via free diffusion under open and absorbing boundary conditions at the beginning and the end states of the process. We conclude that the average successful crossing time is close to but even faster than the mean first-passage time of a freely-diffusing particle, due in part to including time for the free particle”s failures to translocate across.

Thus, our analysis of simulation trajectories suggests that the process of passive transport through the NPC consists of many unsuccessful translocation attempts interrupted by infrequent successful translocation. The timescale of successful translocations, however, is found to be prescribed by the time scale of protein free diffusion. We interpret these observations as a process where constantly rearranging FG-nups open and close transient passages that a protein can take to cross from one side of the NPC to the other at speeds prescribed by free diffusion. Note that this scenario contrasts to a situation where the action of FG-nups is to slow down the effective diffusion of the proteins, through either entanglement or binding, which would manifest itself in a slower than free diffusion timescale of successful crossings.

### Barrier to passive transport as a percolation transition

We prove our open passage model of passive transport by developing a theoretical approach that can predict the rate of passive transport of proteins through the NPC from the analysis of equilibrium fluctuations of the FG-nup mesh alone. For each instantaneous configuration of the mesh sampled by the equilibration trajectories, Fig. 2, and for each spherical probe of radius *R*_*p*_, Supplementary Table 2, we classified each voxel within the volume of the NPC as available if the sphere could be placed at that voxel without clashes with either the mesh, the scaffold, the envelop or the confinement potential, producing a 3D map of internal voids within the NPC volume, Fig. 4a, see methods for a detailed description of the procedure. The 3D void map was converted into a 1D potential occupancy map, *P*_1_(*z*), by splitting the NPC volume into disc segments along the pore axis and calculating the the fraction of available voxels in each segment, Fig. 4b. Trajectory average of the potential occupancy function, Fig. 4c, was converted into an effective PMF, Fig. 4d, through Boltzmann inversion. Gratifyingly, the shape of the resulting PMF reproduces the shape of the PMF extracted directly from brute force simulations, Fig. 2f.

**Fig. 4:**
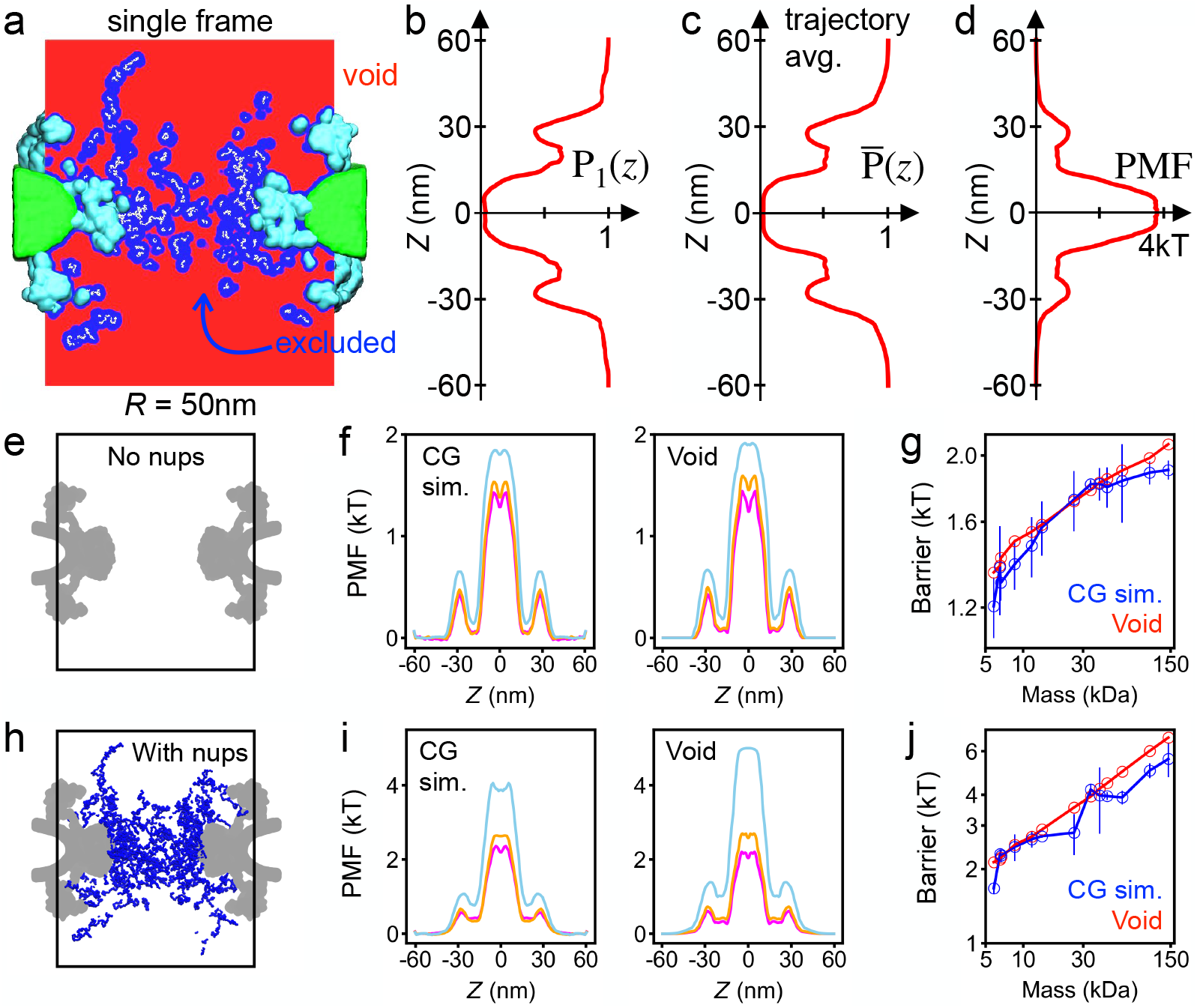
Void model of the translocation barrier. **a** Void analysis map of an instantaneous NPC configuration computed using a spherical probe of 22.4 Å radius. The volume available to accommodate the probe (void) is shown in red, the volume excluded in blue, FG-nups in white, the scaffold in cyan and the lipid bilayer in green. The image shows a 2D section of a 3D map. **b** The fraction of the NPC volume that can accommodate the probe without clashes as a function of the pore axis coordinate. The fraction was computed by splitting the void analysis map into cylindrical segments of 50 nm radius and 0.6 nm height, coaxial with the pore. The data shown were computed for the instantaneous NPC configuration displayed in panel a. **c** Trajectory-averaged probability of accommodating the probe as a function of the pore axis coordinate, 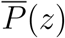, computed by averaging instantaneous void analysis maps over the last 6 ms of the NPC equilibration trajectory, sampled every 1.0 *µ*s. **d** PMF of the spherical probe derived by void analysis. **e** Cut-away view of the NPC system containing no nups, only the scaffold and nuclear envelope potentials (gray). **f** Symmetrized PMF of three protein species (aprotinin, magenta; thioredoxin, orange; and hemoglobin, light blue) derived from brute force CG simulations (left) and of the three spherical probes of approximately the same radius (*R*_p_ =12.75, 15.71 and 30.04 Å) derived by void analysis (right). **g** PMF barrier versus protein mass. Interpolation was used find void analysis PMF barriers for the proteins simulated using the CG method, Supplementary Fig. 8. Lines are guides to the eye. Both axes use logarithmic scale. **h–j** Same as in e–g but for the complete NPC model (including nups).

Further comparison shows that our void analysis method quantitatively reproduces the proteins” PMF. To make such comparison possible, we used interpolation to convert the radius of a spherical probe to the corresponding protein mass, Supplementary Fig. 7, and to find the height of the PMF barrier, Supplementary Fig. 8. In the case of a nup-less NPC, Fig. 4e, the method reproduced fine features of the PMF, including a small dip near the pore midplane (at *z* = 0), Fig. 4f, which is caused by a small widening of the NPC scaffold at the very center of the inner ring. The method also reproduces the absolute height of the barrier, Fig. 4g, without any adjustable parameters. Good quantitative agreement was also observed for the complete NPC model, Fig. 4h–j, and also for simulations carried out under a narrow confinement potential, Supplementary Fig. 9. We note that, for two trajectories of the same length, the data derived from the void analysis method have better statistical sampling because the method characterizes potential occupancy of the entire simulation volume, whereas sampling in a CG simulation is limited to the location of the diffusing protein. The void analysis method also provides a means to estimate the free-energy barrier to passive diffusion of proteins that are too large to obtain good passage statistics from brute force simulations.

Matching the definition of the first-passage used to characterize protein permeation in our CG simulations, Fig. 2h, we compute MFPT by numerically solving the Fokker-Planck equation using our void analysis PMFs and position-dependent diffusion constants, see Methods for detailed description of the procedures. Supplementary Fig. 10 shows the obtained dependence of the MFPT on the void probe radius as well as the interpolation scheme used for direct comparison of the data with the results of the CG simulations. For both nup-less and complete models of the NPC, the MFPTs computed using our Fokker-Planck approach are in excellent agreement with the CG simulation data for both wide, Fig. 5a,b, and narrow, Supplementary Fig. 11a,b, confinement potential case, as well as with the simulation results of a previous study^8^ when scaled by the MFPT of hemoglobin, Supplementary Fig. 12. Note that, in comparison to the Fokker-Planck approach, the CG simulations are expected to systematically underestimate the MFPT of larger proteins because the MFPT distributions are expected to have long tails that are not sampled by the limited-duration CG simulations.

**Fig. 5:**
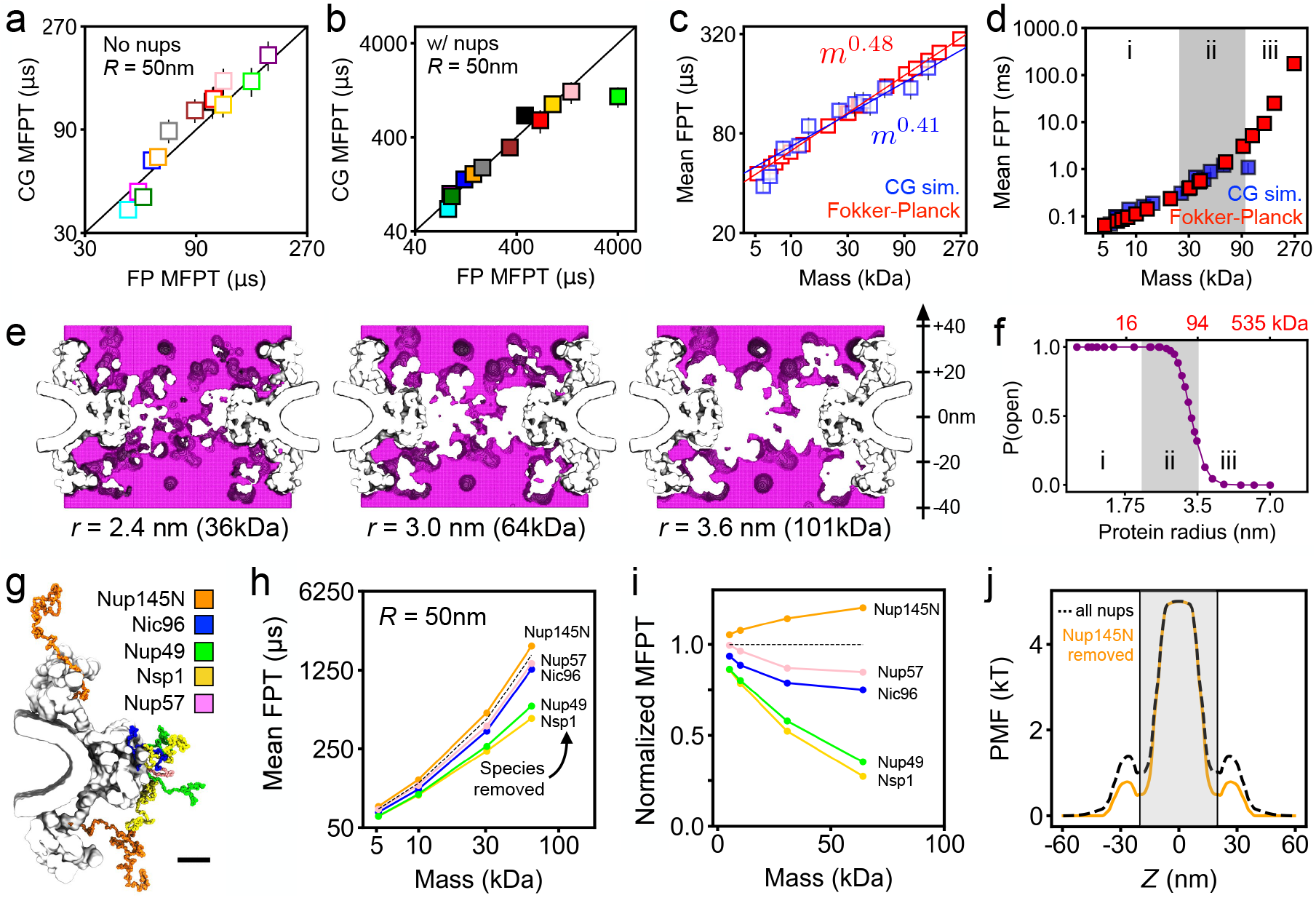
Fokker-Planck model of passive diffusion and the percolation transition. (**a**,**b**) Comparison of the mean first-passage times (MFPT) calculated from CG simulations and using our Fokker-Planck void analysis model for nup-less (panel a) and complete (panel b) NPC models. The black line indicates perfect agreement. Error bars for the CG values indicate standard error. (**c**) MFPT from CG simulations (blue) and using our Fokker-Planck approach (red) as a function of protein molecular mass. Note the logarithmic scale of the axes. Power law fits, and their slopes, are specified in the figure. **d** Same as in panel c but for a complete NPC model, with all FG-nups present. The three regions (i, ii, iii) correspond to power-law, transition and exponential scaling behavior. **e** Connectivity map of an instantaneous NPC configuration computed for three protein probes of different radius (specified under each map). Each map was generated for |*z*| *<* 40 nm. **f** Probability of finding a complete, open path through the NPC versus protein radius and molecular mass (red, top) obtained from the analysis of NPC equilibration trajectories. Note the logarithmic scale of the horizontal axes. The three regions (i, ii, iii) correspond to those in panel f. **g** Location of each FG-nup species in one sixteenth of the CG model. Black scale bar, 10 nm. **h** MFPT versus molecular mass for an NPC model devoid of one FG-nup species (colors) and with all FG-nups present (dashed black line). **i** MFPT for the deletion mutants normalized by the all species present MFPT. **j** PMF of a 64 kDa (*r* = 3.0 nm) protein for all species present NPC (dashed black line) and the Nup145N deletion mutant (orange). The gray shaded region (|*z*| *<* 20 nm) indicates the region used to define the pore length of the central channel. All data in this figure were obtained under a 50 nm radius confinement potential. Interpolation was used to express the results of the Fokker-Planck void analysis model in terms of molecular mass, Supplementary Fig. 10.

In the absence of nups, both approaches yield nearly identical dependences of the MFPT on the protein mass, with a single power law spanning the entire range of protein masses, Fig. 5c and Supplementary Fig. 11c. In the presence of FG-nups, the dependence no longer could be described by a single power law, Fig. 5d and Supplementary Fig. 11d. First, we note that even the largest protein (∼11 nm in diameter) examined above was considerably smaller than narrowest cross section (44 nm in diameter) of the NPC scaffold, although the latter can vary among biological species,^42^ the development stage of an organism^44^ or even tension.^43^ For larger proteins, we indeed observe the nuclear pore to serve as a barrier to passive diffusion; slowing down spontaneous translocation by a factor of 1,000, compared to an empty pore, for the largest protein considered in this study.

To systematically investigate the crossover behavior, we fitted the dependence of the MFPT on the radius using the following expression: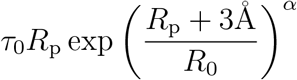, which derives from the transition rate expression *e*^*H*^*/D*, where *H* is the scaled barrier height and the diffusion coefficient *D* scales with the protein radius as 1*/R*_p_. Above, 3Å was added to the protein radius to take into account the physical size of the FG-nups. The fitting yields *τ*_0_ = 3.86 *µ*s/Å, *R*_0_=20.5 Å and *α* = 1.89 for the 50 nm-radius confinement and *τ*_0_ = 1.48 *µ*s/Å, *R*_0_=21.7 Å and *α* = 1.97 for the 25 nm one. When *R*_p_ *< R*_0_, the relation between MFPT and *R*_p_ resembles a power law since the exponential functions could be approximated by two-term Taylor series. The exponential term dominates when log(*τ*_0_*r*) becomes smaller than [(*R*_p_ + 3Å)*/R*_0_]^*α*^, which, in the case of the 50 nm confinement, corresponds to *R*_p_ *>* 35.4 Å and to *R*_p_ *>* 44.9 Å for the 25 nm one. Accordingly, we can divide the dependence of the MPFT on protein mass, Fig. 5d, into a power-law, transition and exponential scaling regions, which suggests a transition from a soft to hard translocation barrier with increasing protein mass.

To determine the physical origin of the crossover behavior, we analyzed connectivity of the transient voids formed within the FG-nup mesh. For each instantaneous mesh configuration, we search the 3D void map for a path crossing the NPC volume (|*z*| *<* 20) that is accessible to a protein of a given radius. To illustrate the procedure, Fig. 5c shows cross sections of three void maps computed for the same mesh configuration and three proteins of different effective radii, i.e., 2.4, 3.0 and 3.6 nm. Upon increasing the protein radius, the path connecting the two sides of the NPC became fragmented as some voxels near the inner ring of the NPC scaffold became inaccessible. We estimated the instantaneous probability to have an open path for a protein of a particular size by performing the above analysis on more than 8,000 FG-nups configurations obtained from the equilibration CG simulations of the NPC model, Fig. 1a. The resulting probability function, Fig. 5d, displays a transition from always having at least one fully-connected path to not having a path at most of the time, a percolation transition. Interestingly, the percolation transition, Fig. 5d, occurs at the same protein mass as where the dependence of the MFPT on protein mass changes from power-law to exponential, Fig. 5b.

We interpret the results of our analysis in a physical model of passive transport where, for small proteins, an open path through an NPC always exists and the protein diffusion is limited by the likelihood of finding this path by diffusion. For larger proteins, the rate of passive transport is also conditioned by small yet finite probability of forming an open path through the FG-nups as a transient fluctuation, resulting in a stronger barrier to translocation that sharply increases with protein mass.

Using our theoretical model, we determined the relative contribution of each FG-nup species to the diffusion barrier. Five additional equilibration trajectories were generated by computationally removing all residues of each of the five FG-nup species, Fig. 5e. The resulting trajectories were analyzed using our theoretical model yielding the dependence of the MFPT on the protein mass for each deletion version of the NPC, Fig. 5f. Normalized by the MFPT observed for the complete NPC model, we find Nsp1 and Nup49 species to contribute the most to the diffusion barrier, Fig. 5g, which could have been expected given their length and the tethering position. The small increase of the MFPT produced by removal of Nup145N is explained by the small change in the PMF which deepened the PMF minimum between the inner and outer rings of the scaffold (near *z* = ±20 nm) without affecting the height of the primary barrier, Fig. 5h. As the location of those PMF minima coincide with the span of the computational domain used to solve the Fokker-Planck equation, removal of Nup145N effectively made the translocation barrier larger, increasing the MFPT.

## Conclusion

We have constructed a structurally accurate computational model of the NPC and carried out CG simulations to find the passive transport through the NPC to be dominated by rare, fast crossings. We have shown that such crossings occur in a free diffusion regime, which suggests a mechanism where the protein transport through the mesh is conditioned by the presence of open paths connecting one side of the NPC to the other. We have shown that to be the case by constructing a theoretical model of passive transport derived from geometrical analysis of the volume available to accommodate the translocating molecule. In that respect, our model is similar to the approach used by Bodrenko and co-workers to described passive diffusion of antibiotics through a membrane channel.^53^ Based solely on the analysis of the free volume, our theoretical model could not only reproduce the transport rates observed in our CG simulations but also allowed us to estimate the transport rates for the proteins that were too large to be characterized through brute force simulations. We find that the average translocation time initially increases proportionally to the protein molecular mass but undergoes a crossover to an exponential dependence for larger masses. By analyzing how the connectivity of the two compartments depends on the protein mass, we associate the crossover in the scaling behavior with a percolation transition, which indicates a qualitative change in the character of the passive transport from being dominated by the protein finding an entrance to a connecting path to the proteins waiting for the connecting path to form. This change in the rate limiting step enables the NPC to function as a protein size filter presenting a soft barrier to diffusion of small proteins and a hard barrier to transport of larger proteins, with the protein size cutoff being determined by the percolation transition.

Our void model of passive transport has several limitations. The model is designed to work for globular proteins that do not exhibit specific binding to FG-nups, which would be the case for the assisted nuclear transport. It is, however, conceivable that introducing one or more binding site along a quasi 1D path through the NPC mesh could increase the rate of transport in comparison to the passive diffusion case, as suggested by previous theoretical work.^23–26^ Although the model assumes the proteins to be spherical, non-spherical shapes can be accommodated in the model at the level of the PMF calculations by modifying the void search protocols. ^54^ The model is not expected to work for partially or fully disordered proteins or other polymer-like molecules (such as mRNA) because polymer entanglements are not accounted for in our Fokker-Planck formulation of the transport. The void analysis approach could be extended to describe a situation where multiple protein species diffuse through the mesh by iterating the void finding procedure with a probabilistic placement of proteins within the voids according to the proteins” PMFs and bulk concentrations, until convergent PMFs profiles are established for all protein species.

Our experimentally-derived CG model sets the stage for further computational exploration of the nuclear transport. We envision applying our model to study passive transport through alternative models of the NPC structure^42,43^ as well as incorporating specific binding in the model to study assisted transport of larger cargoes. Accounting for the chemical reactions that give the nuclear transport its directionality and for the recycling of the transport factors would furnish the first complete physics-based model of nuclear transport.

## Methods

### CG simulations of passive protein transport through the NPC

In our CG model of the NPC, the disordered mesh region is represented using a beads-on-astring model previously developed by the Onck lab^49,50^ whereas the nuclear envelope and the structured protein scaffold are represented using custom grid-based potentials. The proteins diffusing through the NPC are described as rigid bodies. Each component of the model is described in detail below.

#### CG model of the NPC

In accordance with the stoichiometry of the composite NPC model,^46^ the disordered mesh regions consisted of 32 copies of each of the following five species: Nup145N, Nic96, Nsp1, Nup49, Nup57, 160 individual protein chains in total. The amino acid sequence of the disordered domain of each protein species was determined by comparing the full-length protein sequence (UniProt entry G0SAK3 for Nup145N, G0S024 for Nic96, P14907 for Nsp1, G0S4X2 for Nup49 and G0S0R2 for Nup57) to the corresponding sequence of the protein domain resolved in the structure of the protein scaffold.^46^ Thus, the disordered mesh region consisted of 732 (Nup145N), 139 (Nic96), 467 (Nsp1), 245 (Nu49) or 77 (Nup57) amino acids, each being a continuous N-terminal fragment of the corresponding full-length protein. Each amino acid residue of the fragments was represented by one CG bead; the beads were connected into a polymer chain via a harmonic spring potential, see Refs. 49 and 50 for details. We validated our implementation of the CG model by simulating a 10×10 array of Nup62 anchored onto a flat surface and measuring the maximum height of the Nup62 brush, which converged to the same value (12 nm) as in Fig. S1 of Ref. 50. The Cterminal bead of each fragment was harmonically restrained (*k* = 10.0 kcal mol^*−*1^ Å^*−*2^) to the location prescribed by the composite NPC model.^46^ The initial configuration of each fragment was generated as a self-avoiding random walk, with the distance between consecutive beads of 0.38 nm and under conditions that no three consecutive beads of the same fragment were more than 0.8 nm apart, no beads of separate fragments were within 0.8 nm of each other and no beads at least three residues away from the anchor point were within 0.8 nm of the protein scaffold. Two complete sets of 160 FG-nup fragments were generated independently, providing initial conditions for the two replica simulations that utilized the same model of the nuclear envelope and the protein scaffold.

The nuclear envelope was represented using a purely steric, repulsive, potential. The overall shape of the potential was obtained by manually fitting a surface to the center line of the apparent lipid bilayer density seen to surround the NPC in the cryo-ET structure, EMD-3103, ^47^ resulting in the following parametric equation for the surface:

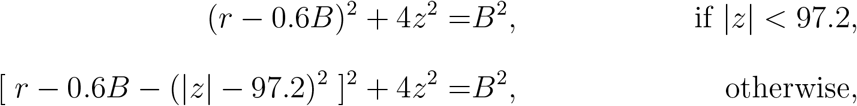

where *z* and *r* are the pore axis and radial coordinate, respectively, in Å, and *B* = 300 Å. The first equation above represents an ellipse, translated away from the origin and rotated about the pore axis. The second equation effectively flattens the curvature of the ellipse so that the lipid bilayer is not raised in the corners of the system. Using a custom Python script, the surface was converted to a triangular mesh consisting of 1,266 faces and an average area of individual triangles of ∼25 nm^2^. The triangular mesh was provided to the `lipidwrapper` tool^55^ along with a square (20 nm on side) patch of a diphytanoyl phosphatidylcholine (DPhPC) membrane, which was replicated and transformed to construct the curved nuclear envelope without gaps. The resulting all-atom model was cut along the *x* and *y* axes at ±60 nm. The all-atom representation was converted to a 120 × 120 × 76 nm^3^ number density map (in units of atoms/Å^3^) at 0.4 nm resolution using the `volmap` tool of VMD;^56^ a spherical gaussian blur was applied to each atom with a standard deviation of 8 times the nominal atomic radius. The 3D density map was applied as a potential to the FG-nup beads through linear interpolation, producing a potential that ranged in values from 0 to 15 kcal/mol.

The structured protein scaffold was represented by 20 grid-based potentials, each describing the interaction of an individual amino acid of a particular type with all amino acids of the scaffold. The potentials were derived by first converting the composite all-atom model of the NPC protein scaffold^46^ to a one-bead-per-residue representation. Following that, the number density of type Y residues in the protein scaffold, *ρ*_Y_, was obtained using the volmap plugin of VMD and with each bead being represented using a normalized gaussian distribution of 1.5 Å width. The scaffold potential for residue type X at position 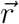 was computed as

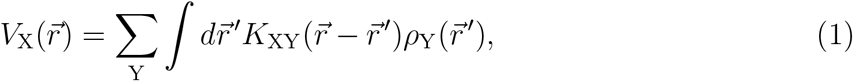

where 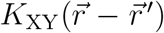 is the potential between beads of type X and Y located at 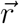 and 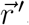, respectively. Eq. 1 was integrated numerically using an FFT-based convolution algorithm using a cubic grid of 0.6 nm resolution.^57^

#### CG simulations of the NPC model

All CG MD simulations were performed using the Atomic Resolution Brownian Dynamics (ARBD) package.^48^ The grid-based potentials representing the nuclear envelope and the protein scaffold were fixed in space. Following Ref. 50, the motions of CG beads representing disordered FG-nup mesh was described using a Langevin dynamics integrator with a timestep of 0.02 ps and the Langevin friction coefficient of 50 ps^*−*1^. The above damping coefficient corresponds to a diffusion coefficient roughly 500 times greater than experimental measurements of individual amino acids.^58^ Hence, we scale the timestep by 500 to obtain an effective timestep of 10 ps. Reporting dynamics using the effective time step is sensible for Rouse-like polymers where hydrodynamic interactions can be neglected. We chose to report our simulation results using the effective time step because experiments have found fully extended intrinsically disordered peptides, similar to FG-nups, to exhibit low internal friction and Rouse-like dynamics. ^59^

The two replica NPC systems were simulated for 7,500 *µ*s each. During the first 1,000 *µ*s, the temperature of the system was reduced from 600 to 298.15 K in 100 evenly spaced steps. The temperature was kept at 298.15 K thereafter using a Langevin thermostat. The systems were simulated under periodic boundary conditions with a unit cell measuring 120 × 120 × 360 nm^3^. Bead–bead and bead–potential forces were calculated every simulation step with a cutoff of 5.0 nm and a pairlist distance of 6.0 nm; the pairlist was updated every 500 steps. The potentials were linearly interpolated to calculate the forces they exerted on beads. The system”s coordinates were recorded every 10,000 steps and used for further analysis.

#### CG model of proteins diffusing through the NPC

Rigid-body models of the 13 globular proteins were generated by converting their all-atom crystallographic structures (PDB IDs 2HIU, 4PTI, 2SPZ, 1UBQ, 1F6M, 1F6S, 1EMA, 2ABH, 1ANF, 6ENL, 4HHB, 1G5Y and 1U8F) to a one-bead-per-residue representation. Thus, each protein rigid body was a collection of CG beads fixed in relative position to each other. Just like a bead in an FG-nup chain, each bead of the rigid body interacted with the beads of the FG-nup mesh or with the potentials representing the protein scaffold or the lipid membrane. At each time step, the forces (and corresponding torques) on the constituent rigid body beads were added up to obtain the net force (and torque) on the rigid body.

The translational and rotational diffusion coefficients of each protein rigid body were obtained from the atomic coordinates of the protein using the HYDROPRO software package.^60^ Only diagonal elements of the diffusion tensor returned by HYDROPRO were used when integrating the equations of motion.

#### CG simulations of proteins diffusion through the NPC

In our simulations of single rigid body protein diffusion through an NPC, the motion of FG-nup was modeled using Langevin dynamics with an effective timestep of 10 ps, consistent with the original formulation of the CG model.^49,50^ The protein diffusion coefficients along and about the protein principal axes were multiplied by a factor of 500 to make the protein diffusion timescale consistent with that of individual amino acids of the FG-nup mesh. The enhanced diffusion coefficient precluded the use of a Langevin dynamics integration scheme for the proteins, which would generate unreasonable persistent path lengths up to 6.2 nm for the largest protein—a significant fraction of the length of the central channel of the NPC. Hence, we update the coordinates of the protein rigid body each timestep using a Brownian dynamics algorithm with an effective timestep of 10 ps so that the direction of the protein motion is uncorrelated between steps.

In addition to the potential representing the NPC scaffold and the membrane envelope, a rigid-body protein was also subject to a confinement potential that was zero within a cylinder centered at the origin. Outside that cylinder, a harmonic potential acted on the protein”s center-of-mass with a spring constant of 0.01 kcal mol^*−*1^Å^*−*2^. The simulations of protein diffusion were performed using confinement potentials of 50 and 25 nm radius, the height of the cylindrical potential was 120 nm in both cases. The forces between the beads of the rigid bodies were calculated every time step with a cutoff of 5 nm and a pairlist distance of 9 nm. The forces exerted on the rigid bodies by the lipid, scaffold and constraint potentials were calculated by linearly interpolating the grid-based potentials at every step.

For each of the 13 globular protein species represented by rigid bodies, two replica systems were simulated differing only by the initial position of the protein, which was positioned the same (50 nm) distance away from the midplane along the pore axis in either direction. Each system was simulated for 6,000 *µ*s (12,000 *µ*s per species). The coordinates of the NPC beads and of the protein rigid body were recorded every 2,500 steps (25 ns). Additional simulations of protein diffusion were performed in the presence of the nuclear envelope and protein scaffold potentials only, i.e., in a model system containing no disordered FG-nup mesh.

### Analysis of CG simulations

#### Nup density maps

The nucleoporin density maps were generated using the final 6,500 *µ*s of both CG simulations, taking into account the 8-fold rotational symmetry of the NPC about the pore axis, and the 2-fold reflection symmetry with respect to the midplane of the nuclear envelope membrane. Each FG-nup bead was assigned a radius of 0.3 nm and a mass of 120 Da. The average 3D density of the FG-nups was calculated using `volmap` plugin of VMD, over a 120 × 120 × 360 nm^3^ volume. The density maps were generated separately for each of the five nup species; the total nup density was calculated by adding together all the density maps for all species.

#### Void Analysis

To identify the volume that can accommodate, without a steric clash, a protein of radius *R*_*p*_ for a given instantaneous configuration of the NPC, we first partitioned the whole system into cubic cells, each cell having a linear dimension equal to 6 Å for our PMF calculations and 1 Å for the connectivity analysis. Each cubic cell voxel, denoted as (*n*_*x*_, *n*_*y*_, *n*_*z*_), was assigned cartesian coordinates located at the voxel”s center.

For a given configuration of the FG-nup mesh, we first determined the distance *R*_max_ from the center of each voxel to the nearest nup particle surface (taking the nup particle”s 3 Å radius into account) or to the nearest void analysis voxel of non-zero scaffold, lipid or constraint potential (as defined in our CG MD simulation of protein transport), whichever was the closest. Each voxel was then classified as accessible for a protein type with radius *R*_*p*_ if *R*_max_ ≥ *R*_*p*_; otherwise it was classified as inaccessible to the protein. We implemented our algorithm using a cell linked-lists data structure to store the positions of FG-nups particles, achieving *O*(*N*) computational complexity, where *N* is the total number of voxels.

#### Connectivity Analysis

Results from the aforementioned void analysis were applied to perform connectivity analysis, in which we query the existence of a continuous path through the FG-nup mesh for each configuration. A layer of voxels above (at *z* = +20 nm) and below (at *z* = −20 nm) of the NG-nup mesh were defined as the source and sink regions, respectively, for the continuous path search. Voxels between the source and the sink were identified as available (or not) according to the void analysis procedure. After identifying every void voxel for a given protein size, we used a union-find with path compression algorithm ^61^ to partition all accessible voxels into several disjoint sets, where any two neighboring accessible voxels were assigned to the same set. An open path crossing a given configuration of the FG-nup mesh was determined to exist in any of the above sets contained both source and sink voxels.

### Theoretical model of passive protein transport through the NPC

#### Fokker-Planck formulation

Diffusion of a Brownian particle in a potential 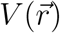 is described within the framework of the Smoluchowski equation

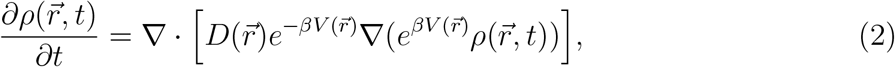

where 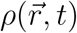 is the Brownian particle density at position 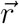 and time *t*, and 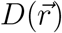 is the position-dependent diffusion coefficient. The geometry of the NPC allows the Smoluchowski equation to be approximated by a one-dimensional differential equation, where the position variable, *z*, is defined along the pore axis.

To solve the equation numerically, we discretize the coordinate space using *N* + 2 points, each separated by distance *d* = 0.5 Å from its neighbor. Each point has a coordinate *z*_*n*_ = *z*_0_ + *nd* with the two boundary points being at *z*_0_ = −600 Å and *z*_*N*+1_ = 200 Å. Following the approach described in Refs. 62 and 63, we rewrite Eq. 2 as

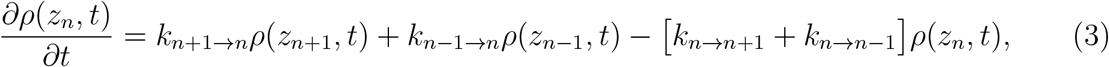

where *k*_*n→m*_ is the transition rate from *z*_*n*_ to *z*_*m*_:

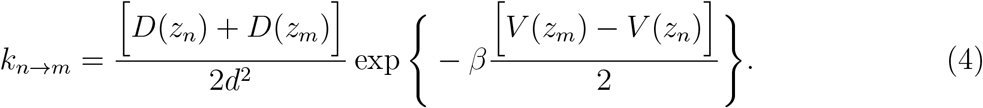

Given the density at time *t*_*i*_, we solved for *ρ* a short time, Δ*t* = 10 fs, later using the Euler method,

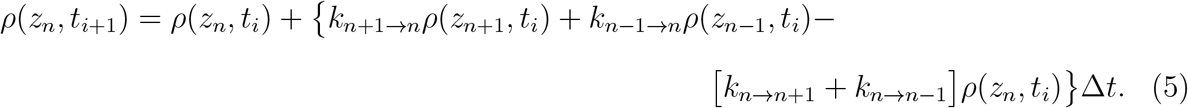

Our choice for Δ*t* = 10 fs satisfies 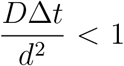 even for the smallest (most diffusive) protein, ensuring convergence of the Euler method. To solve the first-passage time problem, we set an absorption boundary condition at *z*_*N*+1_ and a reflecting boundary condition at *z*_0_:

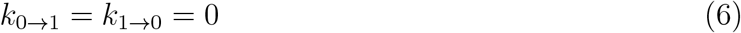

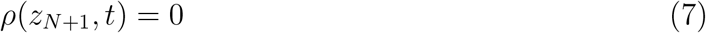

and the initial condition *ρ*(*z*_*n*_, *t*_0_) = *δ*(*z*_*n*_ − *z*^*′*^) assuming the particle starts at *z*^*′*^. In our calculation, *z*^*′*^ is set to −200 Å. The first-passage time (FPT) distribution *g*(*t*) is

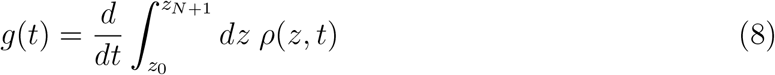

and the mean first passage time (MFPT, denoted *τ*) is then

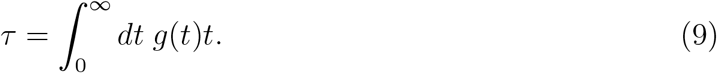

We approximate Eq. 8 and 9 using a finite difference/trapezoidal rule integration scheme.

#### Calculation of PMF from void analysis

To estimate the MFPT from the Smoluchowski equation, the PMF 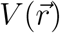 needs to be known *a priori*. Running MD simulations to estimate the PMF for very large proteins is not practical because the slow crossing time would limit sampling. Here we derived an algorithm to estimate the PMF without simulating actual protein translocation.

Neglecting rotational degrees of freedom, the partition function of a system containing FG-nups and one translocating protein can be written,

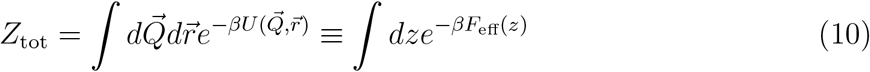

where 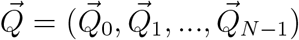 is the vector of the total coordinates for *N* FG-nup particles,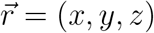 is coordinate of the protein, 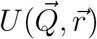 is the total potential energy of the system and *F*_eff_(*z*) is the effective one-dimensional PMF that depends only on the *z* coordinate of the protein. We rewrite 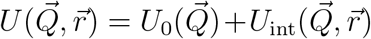, where 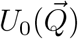 represents the interaction between FG-nup, and 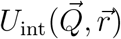 represents the interaction between the protein and the FG-nups. Hence, we rewrite Eq. 10 as

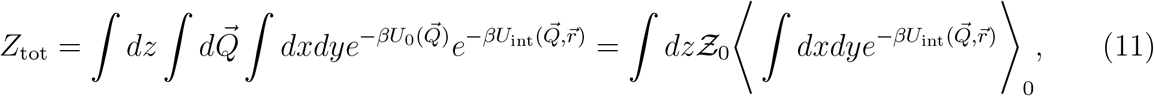

where ⟨…⟩_0_ denotes the ensemble average over the configurations where no protein was present and 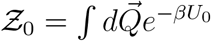 is a normalization constant. Combining Eqs. 10 and 11, we can write

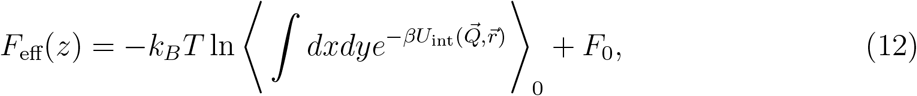

where *F*_0_ comes from the normalization constant. We note that Eq. 12 is well-known as the foundation for free energy perturbation. ^64,65^ Based on Eq. 12, to obtain *F*_eff_(*z*), one can simply generate a canonical ensemble for the system containing only FG-nup particles, and then evaluate the interaction energy for each configuration of the ensemble. The generation of the FG-nup ensemble can be done using brute force Monte Carlo or MD simulations at constant temperature, which is much more efficient than simulating proteins translocating through the NPC. Our derivation was under the assumption that there is only one copy of the protein, however the resulting expression is valid for any dilute system where protein–protein interactions are negligible.

We further simplify Eq. 11 by approximating 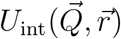 with only steric interaction,

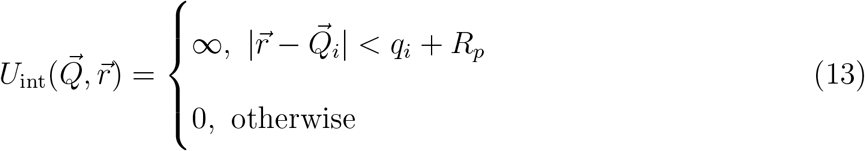

where *q*_*i*_ is the radius of the FG nup particle *i*, and *R*_p_ is the radius of a spherical probe used for the void analysis, Supplementary Table 2.

Using Eq. 13, we express the partition function, *Z*_tot_, and the effective PMF, *F*_eff_(*z*), in terms of the cross section area A(*z*) available for a protein of radius *R*_p_ at pore axis coordinate *z*:

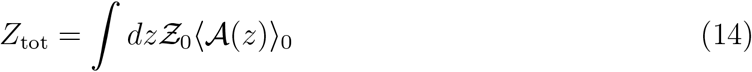

And

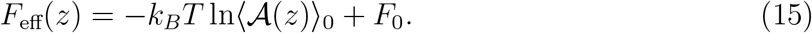

To efficiently evaluate Eq. 15 for proteins of arbitrary size, we first obtained an ensemble of protein-free FG-nup conformations by sampling instantaneous configurations from the 14 ms CG equilibration of the NPC system every microsecond. For each conformation, the FG-nup coordinates were binned into cells of a 3D grid with side-length *l*_*c*_ = 6 Å. For a given protein size and a nup conformation, we then evaluated the histogram *h*(*z*_*i*_) for the number of cells available to the protein at height *z*_*i*_ using the void analysis algorithm described above.

The histogram was then averaged over the ensemble of FG-nups, yielding the effective PMF

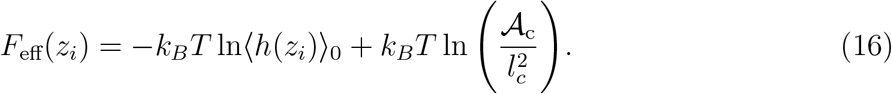

where *𝒜* _c_ is the cross section of the cylindrical restraint potential used in our CG simulations. In the above expression, the second term ensures the PMF is zero far away from the pore. We separately estimated the PMF from the first and second replica trajectories of the FG-nups and found a negligible difference between them. The maxima of the PMFs differed by only 0.6% for a protein of radius 52 Å, the largest radius considered in this study, suggesting all PMF calculations are converged.

Our expression for the PMF, Eq. 15, bears resemblance to the Fick-Jacobs equation describing diffusion of a particle through an entropic barrier.^66^ Following the heuristic argument of Ref. 67, the reduction in the configuration space not only produces the entropic barrier but also has an effect on the diffusion coefficient. Hence, we approximate the diffusion coefficient *D*(*z*) at cross section *𝒜* (*z*) to be

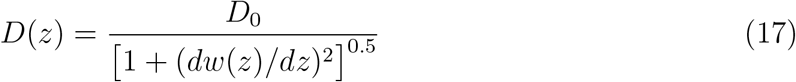

where the effective half width of the channel, *w*(*z*), is defined as

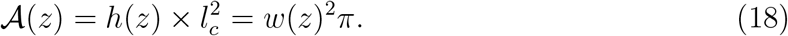

Using the expression for the effective PMF, Eq. 15, and for the position-dependent diffusion coefficient, Eq. 17, we numerically solved Eq. 2 for *ρ*(*z, t*), *g*(*t*) and MFPT *τ* using a custom C code.

The source code and the application examples for the void analysis method and numerical solution of the 1D Fokker-Planck equation are available at https://gitlab.engr.illinois.edu/tbgl/pubdata/void_analysis and https://gitlab.engr.illinois.edu/tbgl/pubdata/fokker_planck_1d, respectively.

## Acknowledgements

This work was supported by the Center for the Physics of Living Cells through the National Science Foundation grant PHY-1430124 and by the National Institutes of Health through grants R01-GM137015 and P41-GM104601. The supercomputer time was provided by the Extreme Science and Engineering Discovery Environment (allocation MCA05S028), the Blue Waters petascale supercomputer system (UIUC), and Leadership Resource Allocation MCB20012 on Frontera at the Texas Advanced Computing Center. Frontera is made possible by National Science Foundation award OAC-1818253. The authors thank Drs. Patrick Onck and Ali Ghavami for sharing files that described potentials of their coarsegrained model. A.A. would like to thank Dr. Cees Dekker for the invitation to join the nuclear transport field.

## Author contributions

D.W. and H-Y.C. contributed equally as co-first authors of the study. D.W. and A.A. conceived the study. D.W and C.M. built the experimentally-derived NPC model. D.W. performed coarse-grained simulations. H.Y.-C. developed and performed void analysis and Fokker-Planck calculations. C.M. and A.A. assisted with analysis done by D.W. and H.-Y.C. All authors contributed to writing and editing the manuscript.

