## Supplementary Materials for "Percolation transition determines protein size limit for passive transport through the nuclear pore complex"

†*Department of Physics,*

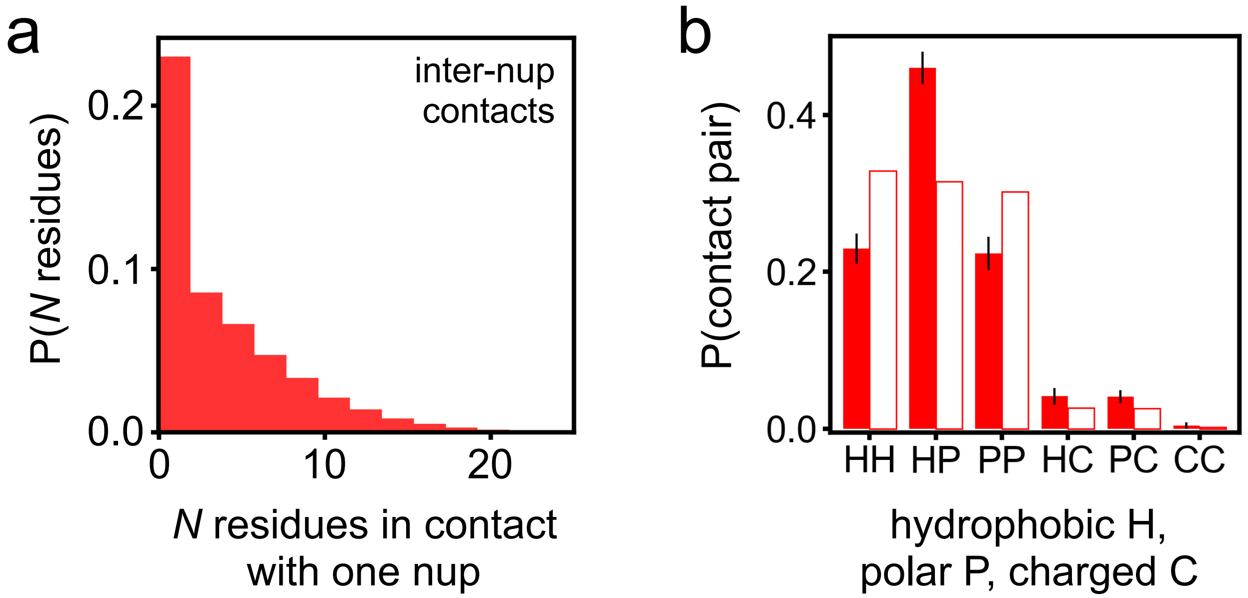


**Supplementary Fig. 1: Contacts between FG-nups. a** The probability of an FG-nup forming contacts with $N$ residues of any other FG-nup. **b** The probability of a pair of residue types to form an inter-chain contact. The residues were categorized as either hydrophobic, H (Ala, Val, Ile, Leu, Met, Phe, Tyr, Trp, Sec, Gly, Pro), polar, P (Ser, Thr, Asn, Gln, His, Cys), or charged, C (Arg, Lys, Asp, Glu). The contact probability extracted directly from CG simulations is shown using filled bars. For reference, we plot the fraction of pair contacts for a random mixture of residues of the same composition as the FG-nup mesh, i.e., $F_{\mathrm{X,Y}} = P_{\mathrm{X}}P_{\mathrm{Y}}/(P_{\mathrm{H}}P_{\mathrm{H}} + P_{\mathrm{H}}P_{\mathrm{P}} + P_{\mathrm{H}}P_{\mathrm{C}} + P_{\mathrm{P}}P_{\mathrm{P}} + P_{\mathrm{P}}P_{\mathrm{C}} + P_{\mathrm{C}}P_{\mathrm{C}})$, where the fraction of residues of type X in the FG-nup mesh $P_{\mathrm{X}} = N_{\mathrm{X}}/(\Sigma_{\mathrm{H,P,C}}N_i)$, $N_{\mathrm{X}}$ is the number of residues of type X in the mesh and X and Y denote either H, P or C. The contact analysis was performed on the final 6,500 $\mu$s fragments of the two CG equilibration trajectories, sampled every 0.1 $\mu$s. A contact was determined to exist when the distance between two amino acid beads from different nups was less than 0.8 nm.

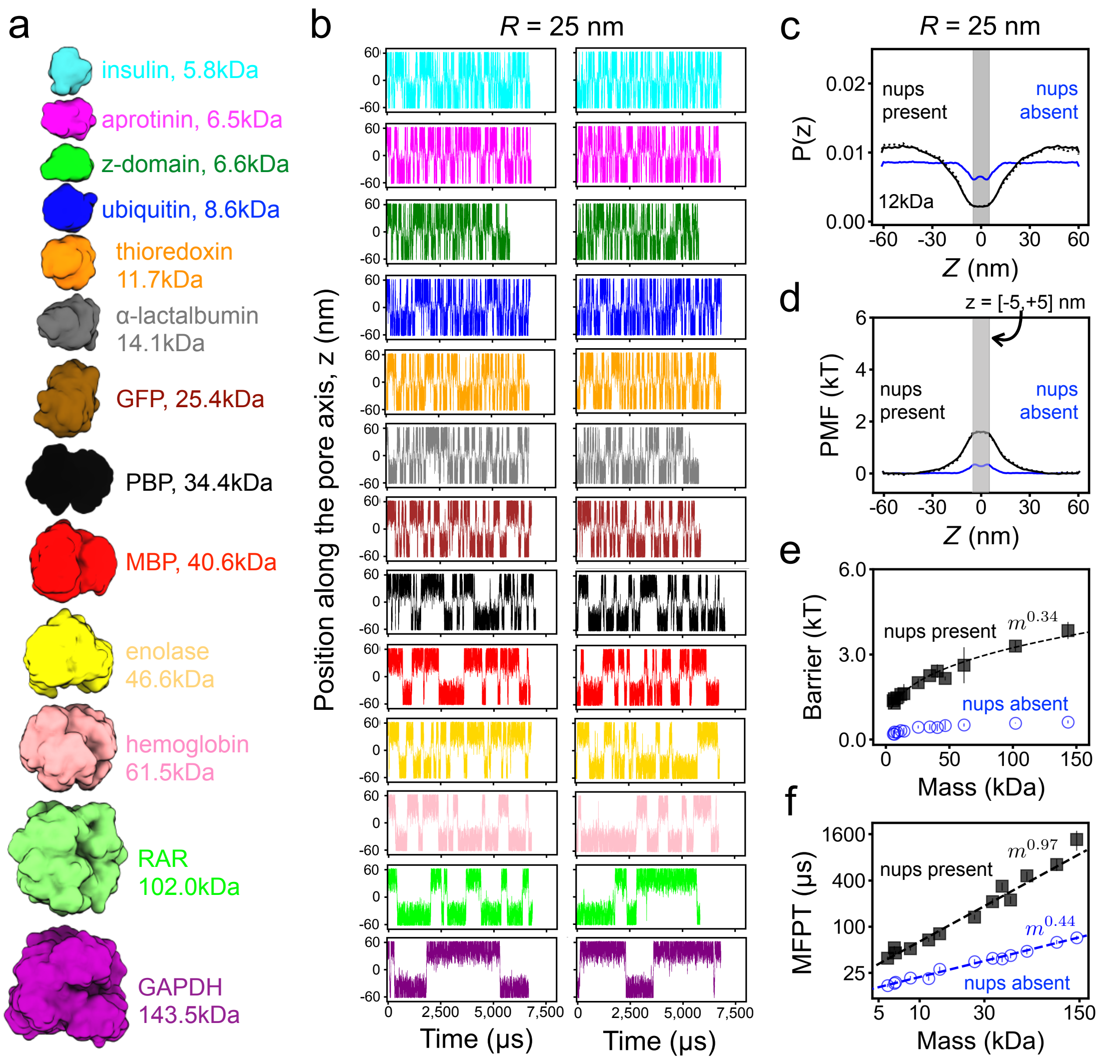


**Supplementary Fig. 2: Passive diffusion across the NPC. a** To-scale structural representation of all proteins used for CG simulations of passive diffusion. **b** Center-of-mass $z$ coordinate of each protein (colors defined in panel A) versus simulation time. The simulations traces in the two columns differ by the initial placement of the protein. The simulation were performed in the presence of a 25 nm-radius confinement potential. **c** Normalized distribution of the CoM $z$ coordinate of thioredoxin. The black dotted and solid lines shows the distribution extracted directly from the simulations and the symmetrized distribution, respectively. The blue line shows a symmetrized distribution for the simulation carried out in the absence of the FG-nup mesh. **d** Potential of mean force (PMF) for thioredoxin transport across the NPC. A PMF barrier is defined as the average value within $|z| < 5$ nm. **e** PMF barrier versus protein molecular mass determined from CG simulations of protein diffusion through our complete NPC model (black squares) and the model missing all FG-nups (blue circles). Error bars are defined as the average point-by-point difference of the unsymmetrized PMF values from $-50 < z < 0$ nm and $0 < z < 50$ nm intervals. Lines show power law fits to the data. **f** Mean first-passage time (MFPT) versus protein molecular mass. Note the logarithmic scale of both axes. Power-law fits are shown as dashed lines in panels e & f. In panel f only, the power-law fit to the "nups present" data included proteins up to 62 kDa (hemoglobin).

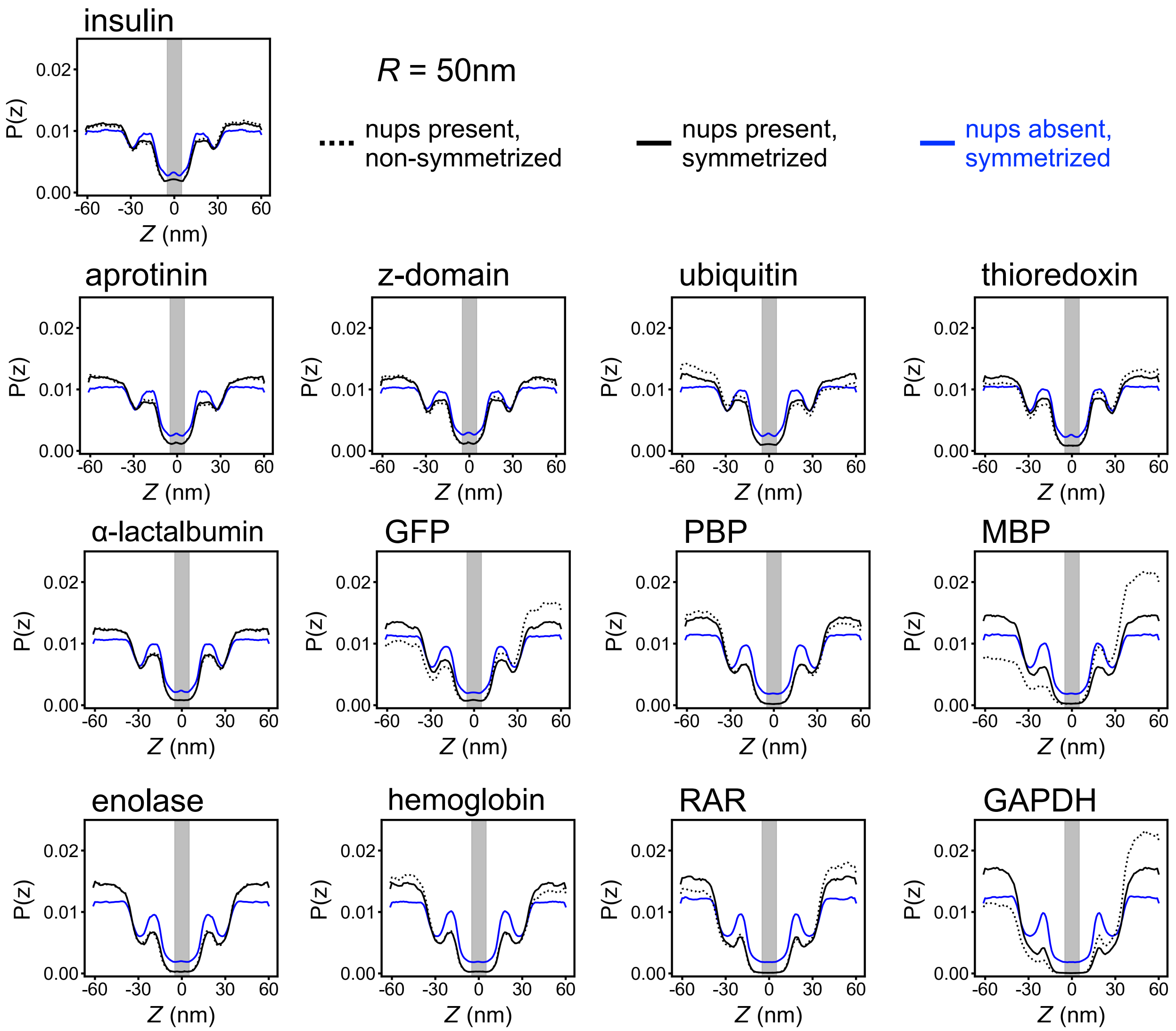


**Supplementary Fig. 3: Protein localization in CG simulations of passive diffusion.** Each panel shows a normalized distribution of the protein's CoM $z$ coordinate. The black dotted lines show the distributions extracted directly from the CG simulations whereas the black solid lines show the same distributions symmetrized with respect to $z$=0. The blue lines show symmetrized distributions for the simulation carried out in the absence of the FG-nup mesh. All data derive from the simulations carried out under a 50 nm-radius confinement potential.

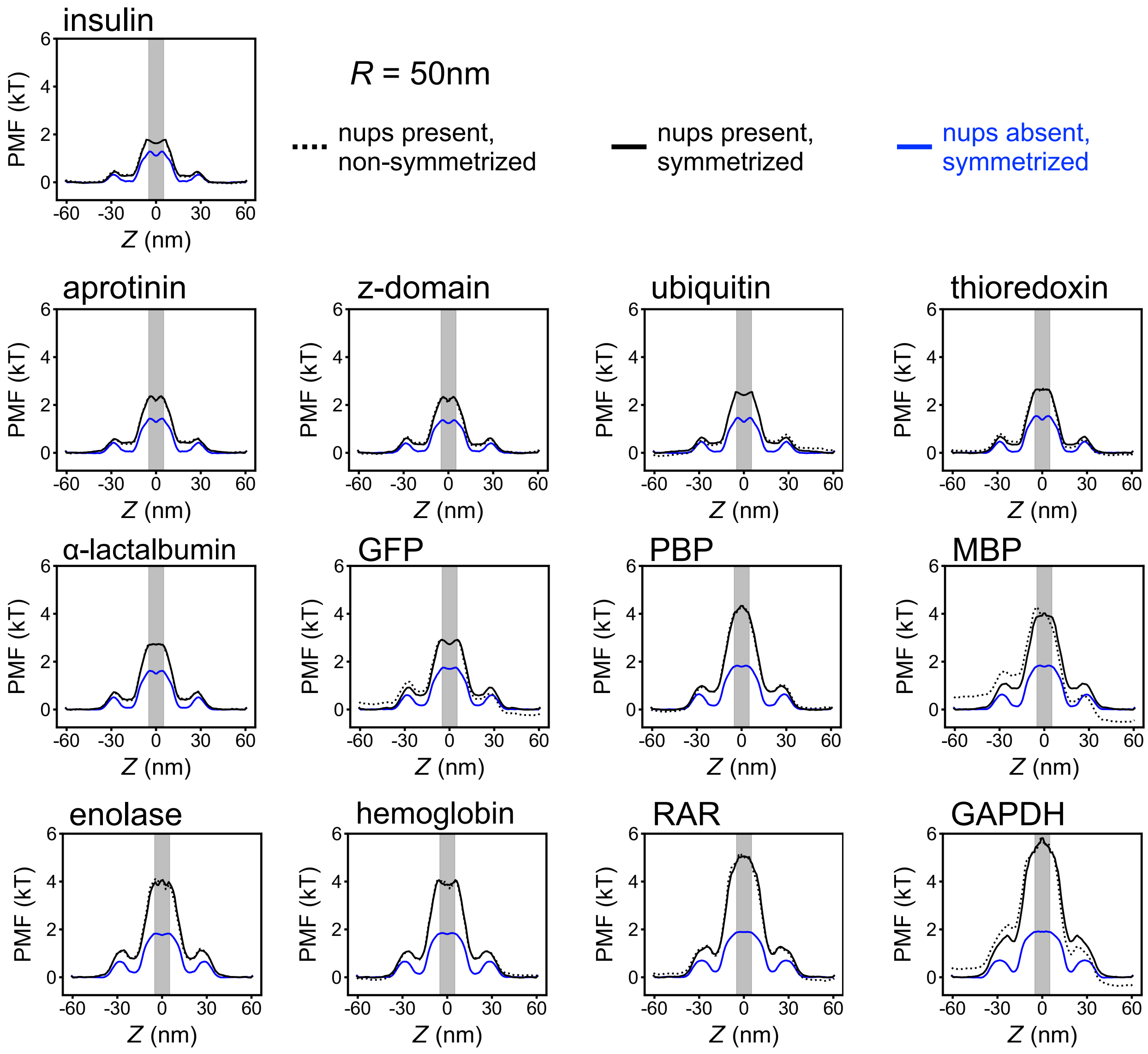


**Supplementary Fig. 4: Potentials of mean force for thirteen protein species extracted from CG simulations of passive diffusion.** The black dotted lines show the PMFs obtained through Boltzmann inversion of the non-symmetrized CoM distributinos, Supplementary Fig. 3. The solid black lines show the same PMFs symmetrized with respect to $z = 0$. The blue lines show symmetrized PMFs obtained from the simulation carried out in the absence of the FG-nup mesh. The grey rectangles illustrate our definition of a PMF barrier. All data derive from the simulations carried out under a 50 nm-radius confinement potential.

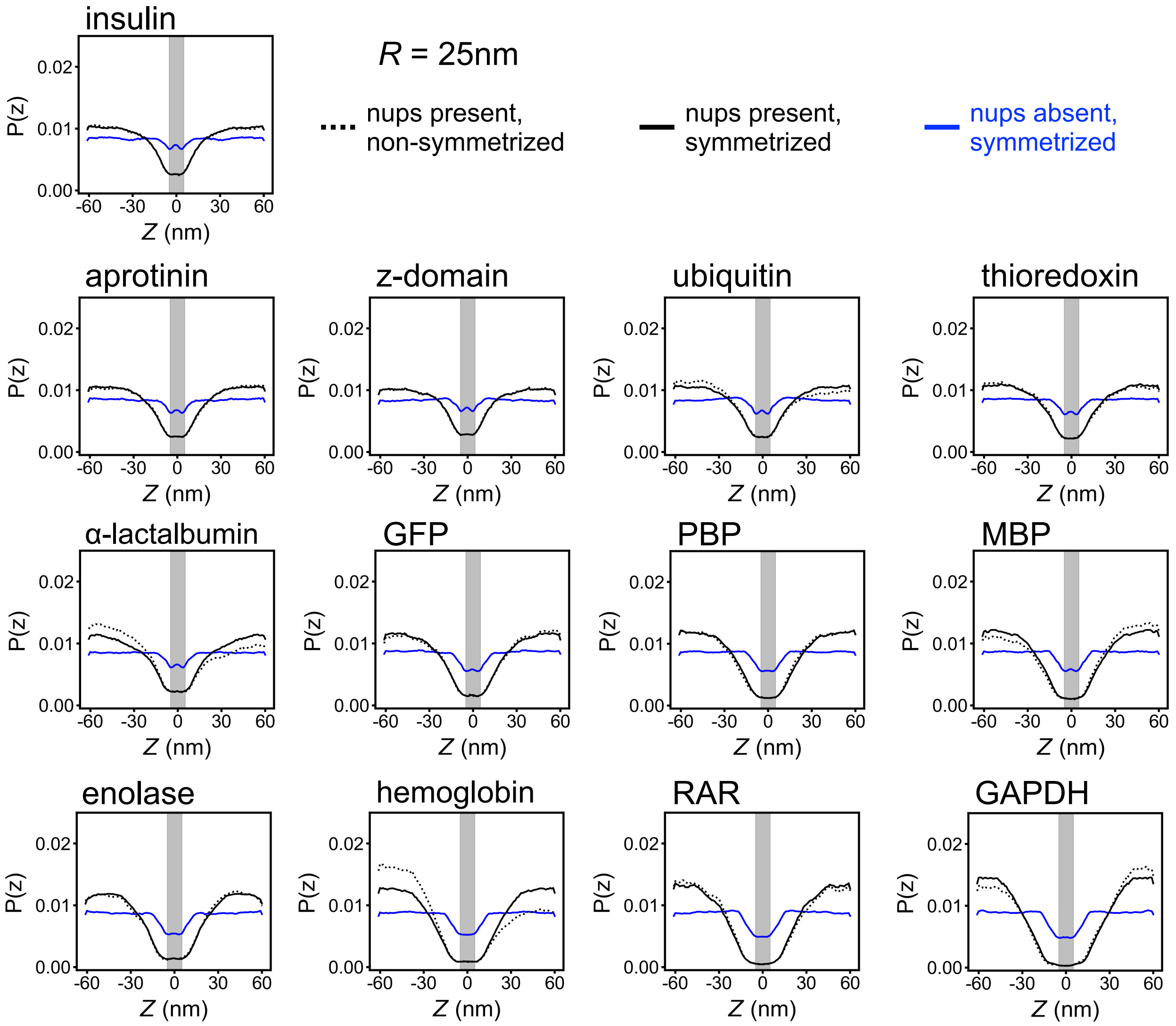


**Supplementary Fig. 5: Protein localization in CG simulations of passive diffusion.** Each panel shows a normalized distribution of the protein's CoM $z$ coordinate. The black dotted lines show the distributions extracted directly from the CG simulations whereas the black solid lines show the same distributions symmetrized with respect to $z$=0. The blue lines show symmetrized distributions for the simulation carried out in the absence of the FG-nup mesh. All data derive from the simulations carried out under a 25 nm-radius confinement potential.

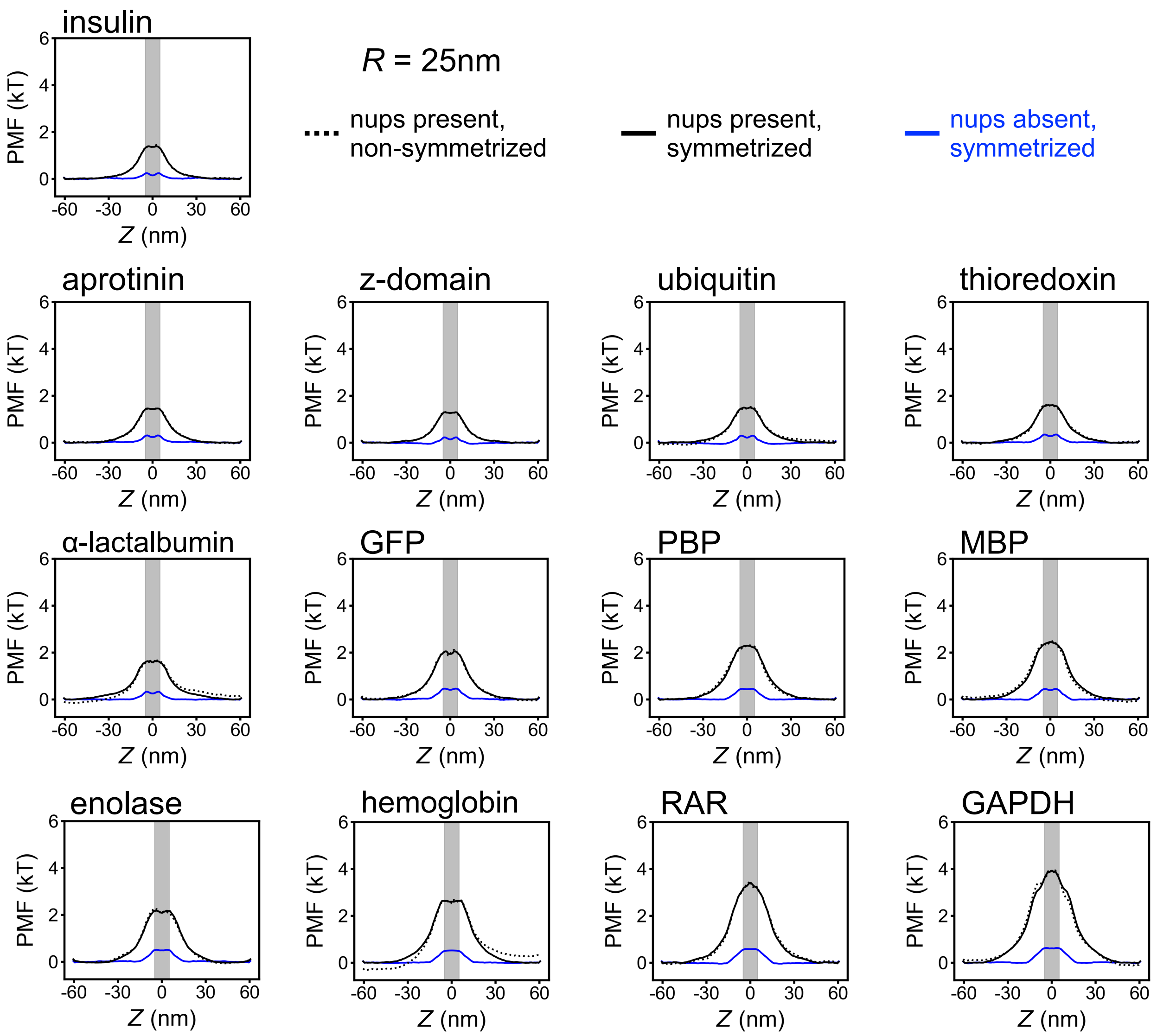


**Supplementary Fig. 6: Potentials of mean force for thirteen protein species extracted from CG simulations of passive diffusion.** The black dotted lines show the PMFs obtained through Boltzmann inversion of the non-symmetrized CoM distributinos, Supplementary Fig. 5. The solid black lines show the same PMFs symmetrized with respect to $z = 0$. The blue lines show symmetrized PMFs obtained from the simulation carried out in the absence of the FG-nup mesh. The grey rectangles illustrate our definition of a PMF barrier. All data derive from the simulations carried out under a 25 nm-radius confinement potential.

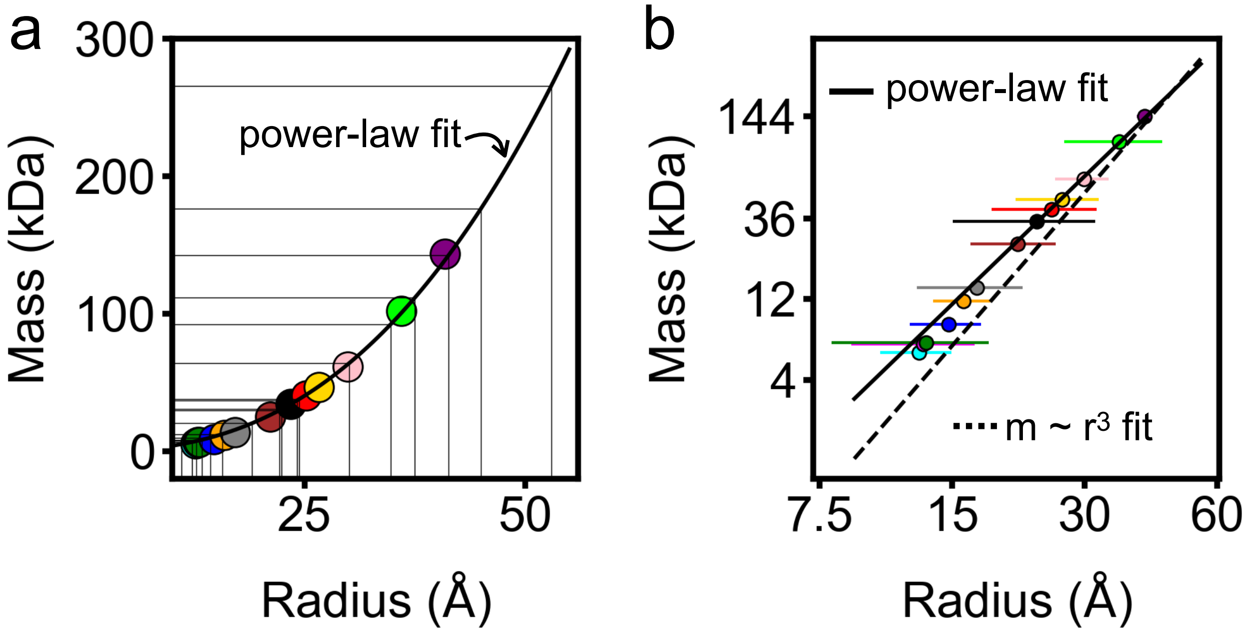


**Supplementary Fig. 7: Molecular mass versus geometric radius of the proteins considered in this work.** Each colored circle represent one protein species simulated using our CG approach. Panels a and b show the same data using linear (panel a) and logarithmic (panel b) scale for the axes. A radius of a protein was computed by matching the protein's moments of inertia with those of an ellipsoid and then approximating the ellipsoid with a sphere of the same volume. The thick solid line shows the best power-law fit to the data, $\sim r^{2.51}$. The dashed line in panel b also shows an $\sim r^3$ fit for comparison. While one would intuitively expect the protein mass to scale as a cube of the protein radius, we empirically found a slower-growing dependence, which we associate with the presence of voids within the proteins. We used this empirical dependence to associate the radius of a spherical probe with a protein mass. Thus, the thin vertical lines represent the radii of the spherical probes used for the void analysis. The thin horizontal lines indicate the equivalent molecular mass for each spherical probe estimated using the best power-law fit. The raw data along with a mathematical formulation of the geometric radius are provided in Supplementary Table 1.

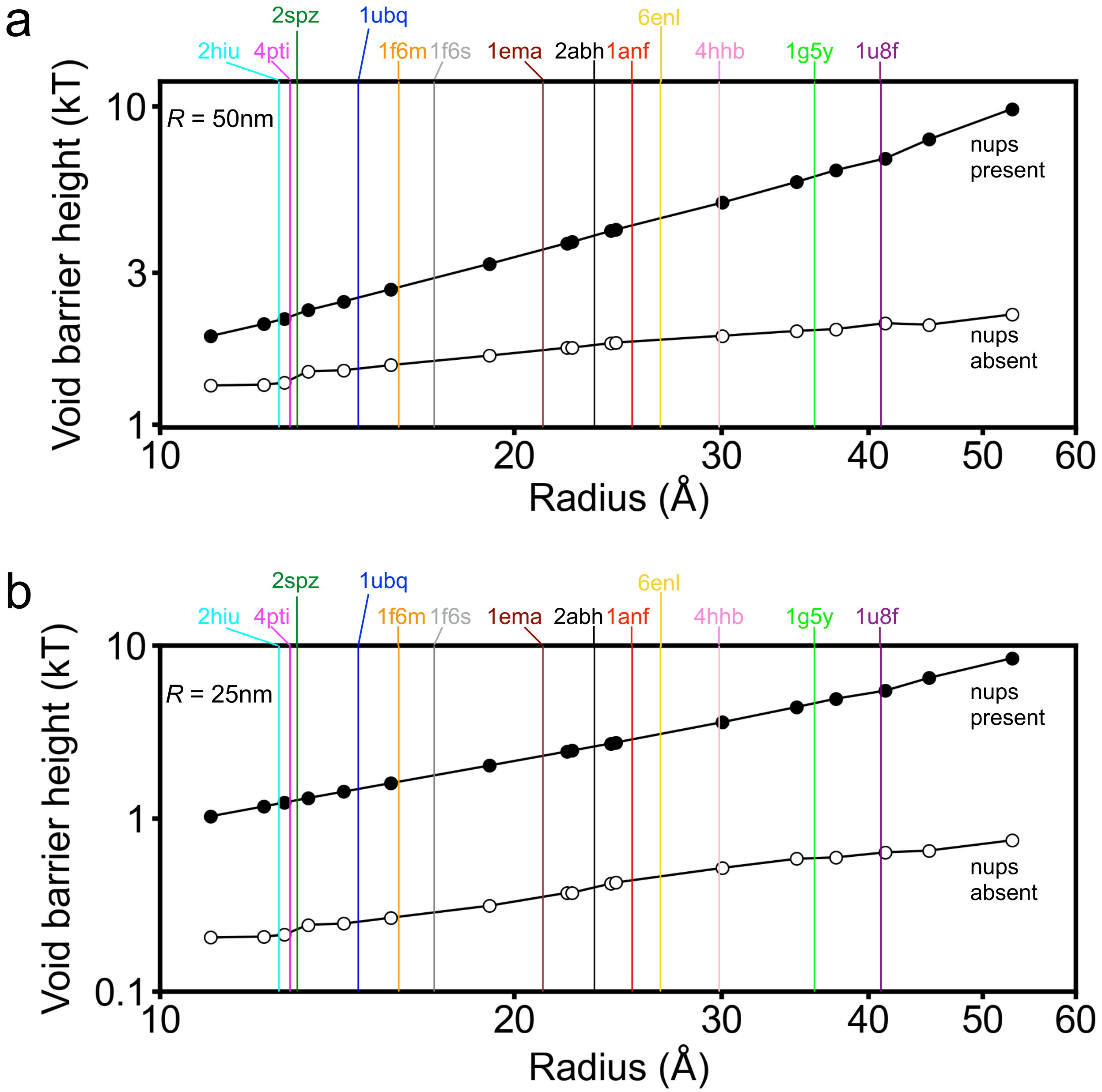


**Supplementary Fig. 8: PMF barrier from void analysis of the FG-nup mesh. a** Height of the PMF barrier versus the radius of the spherical probe used for the void analysis calculations of the translocation PMF. Filled and open circles indicate data obtained for a complete NPC model and a model missing the FG-nup mesh, respectively. Vertical lines indicate geometric radii of the proteins used in our CG simulations; each line is annotated with a corresponding PDB ID. The void analysis value of the PMF barrier for each of the thirteen proteins was determined by interpolation of the void analysis data. Note the logarithmic scale of both axes. These data were obtained assuming a 50 nm-radius confinement potential. **b** Same as in panel a, but for a 25 nm-radius confinement potential.

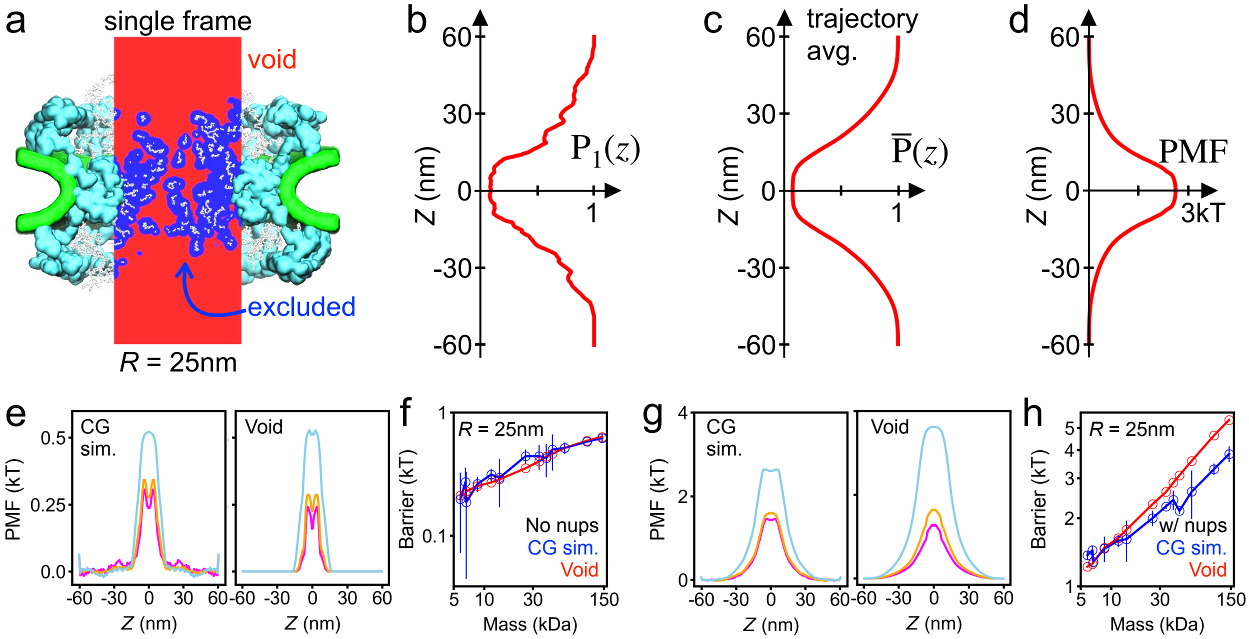


**Supplementary Fig. 9: Void model of the translocation barrier.** **a** Void analysis map of an instantaneous NPC configuration computed using a spherical probe of 22.4 Å radius. The volume available to accommodate the probe (void) is shown in red, the volume excluded in blue, FG-nups in white, the scaffold in cyan and the lipid bilayer in green. The image shows a 2D section of a 3D map. **b** The fraction of the NPC volume that can accommodate the probe without clashes as a function of the pore axis coordinate. The fraction was computed by splitting the void analysis map into cylindrical segments of 25 nm radius and 0.6 nm height, coaxial with the pore. The data shown were computed for the instantaneous NPC configuration displayed in panel a. **c** Trajectory-averaged probability of accommodating the probe as a function of the pore axis coordinate, $\overline{P}(z)$, computed by averaging instantaneous void analysis maps over the last 6 ms of the NPC equilibration trajectory, sampled every 1.0 $\mu$s. **d** PMF of the spherical probe derived by void analysis. **e** Symmetrized PMF of three protein species (aprotinin, magenta; thioredoxin, orange; and hemoglobin, light blue) derived from brute force CG simulations (left) and of the three spherical probes of approximately the same radius ($R_\mathrm{p}$ = 12.75, 15.71 and 30.04 Å) derived by void analysis (right). **f** PMF barrier versus protein mass. Interpolation was used find void analysis PMF barriers for the proteins simulated using the CG method, Supplementary Fig. 8. Lines are guides to the eye. Both axes use logarithmic scale. **g,h** Same as in e,f but for the complete NPC model (including nups).

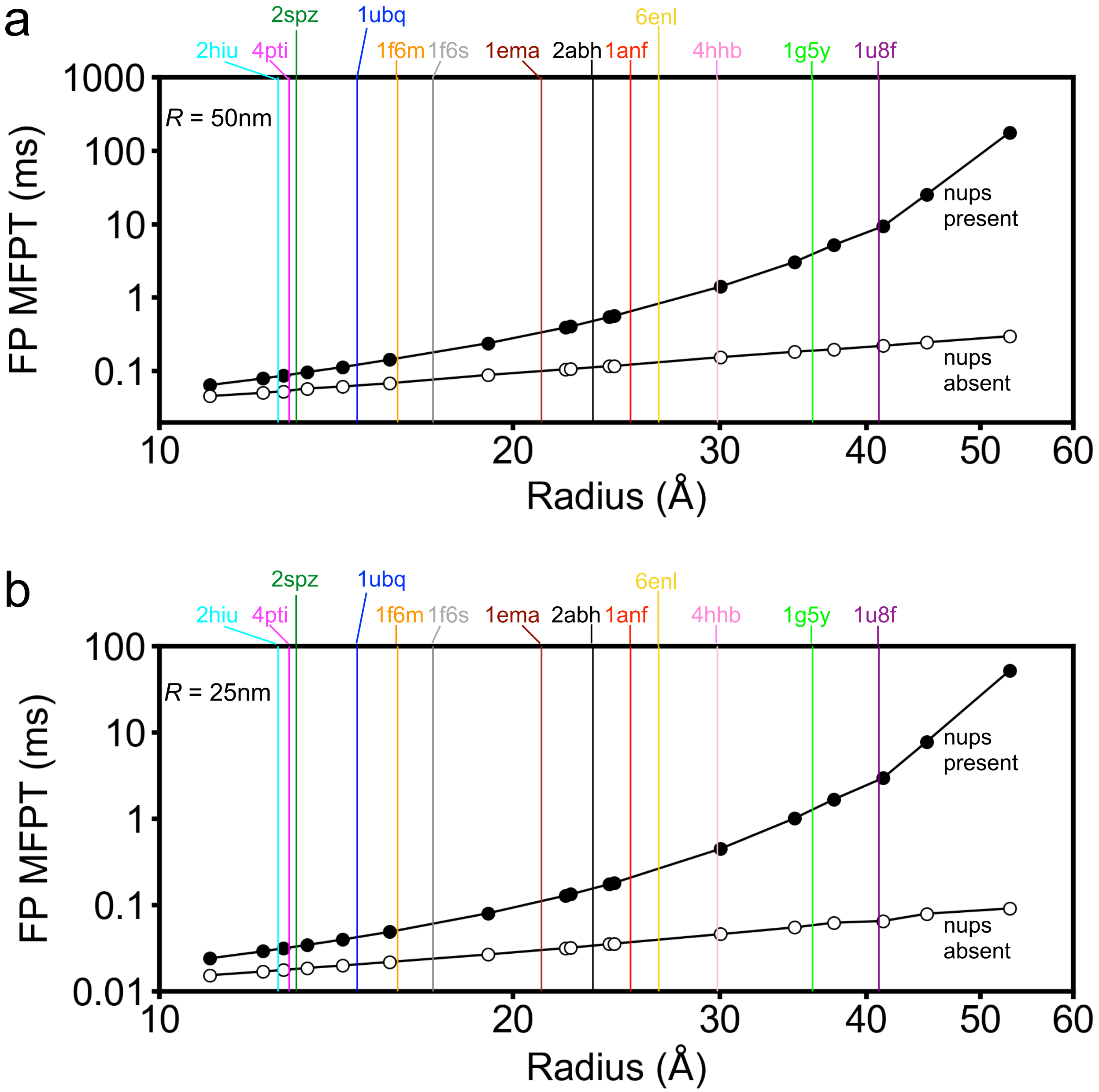


**Supplementary Fig. 10: Mean first-passage time from the 1D Fokker-Planck model.** (A) Mean first-passage time versus the radius of the spherical probe used for the void analysis/Fokker-Planck calculations. Filled and open circles indicate data obtained for a complete NPC model and a model missing the FG-nup mesh, respectively. Vertical lines indicate geometric radii of the proteins used in our CG simulations; each line is annotated with a corresponding PDB ID. The Fokker-Planck value of the mean first-passage time for each of the thirteen proteins was determined by interpolation of the Fokker-Planck data. Note the logarithmic scale of both axes. These data were obtained assuming a 50 nm-radius confinement potential. (B) Same as in panel A, but for a 25 nm-radius confinement potential.

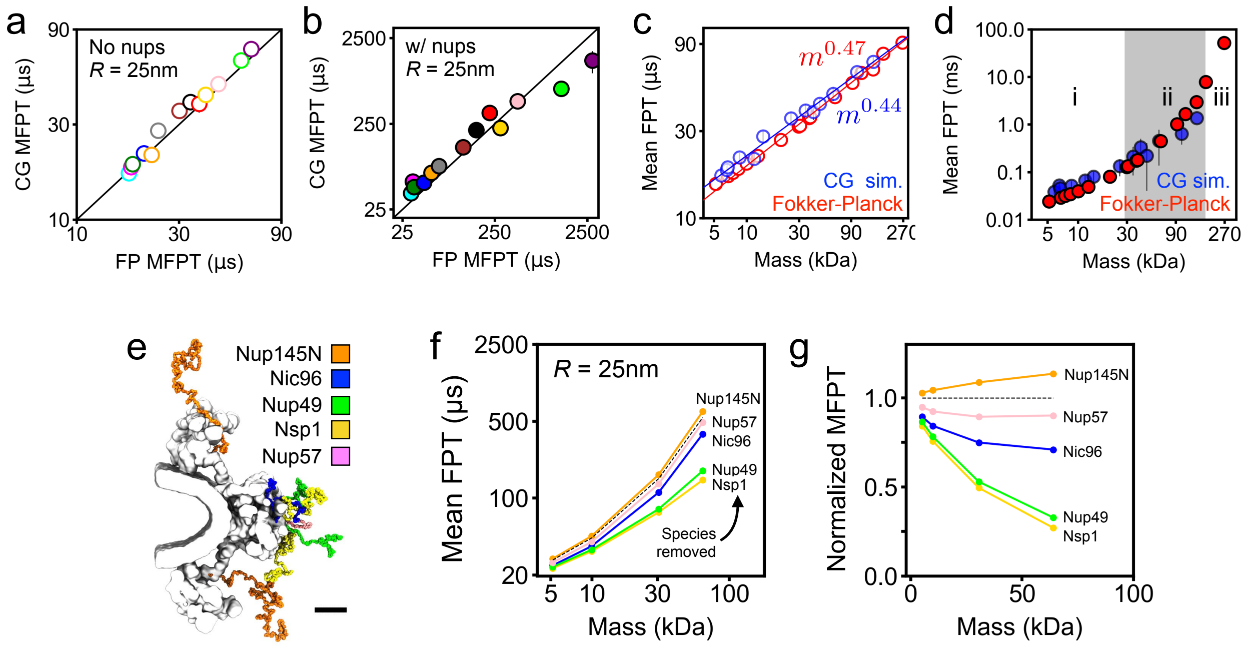


**Supplementary Fig. 11: Fokker-Planck model of passive diffusion through the NPC. a,b** Comparison of the mean first-passage times (MFPT) calculated from CG simulations and using our Fokker-Planck void analysis model for nup-less (panel a) and complete (panel b) NPC models. The black line indicates perfect agreement. Error bars for the CG values indicate standard error. **c** MFPT from CG simulations (blue) and using our Fokker-Planck approach (red) as a function of protein molecular mass. Note the logarithmic scale of the axes. Power law fits, and their slopes, are specified in the figure. **d** Same as in panel c but for a complete NPC model, with all FG-nups present. The three regions (i, ii, iii) correspond to power-law, transition and exponential scaling behavior. **e** Location of each FG-nup species in one sixteenth of the CG model. Black scale bar, 10 nm. **f** MFPT versus molecular mass for an NPC model devoid of one FG-nup species (colors) and with all FG-nups present (dashed black line). **g** MFPT for the deletion mutants normalized by the all species present MFPT. All data in this figure were obtained under a 25 nm radius confinement potential. Interpolation was used to express the results of the Fokker-Planck void analysis model in terms of molecular mass, Supplementary Fig. 10.

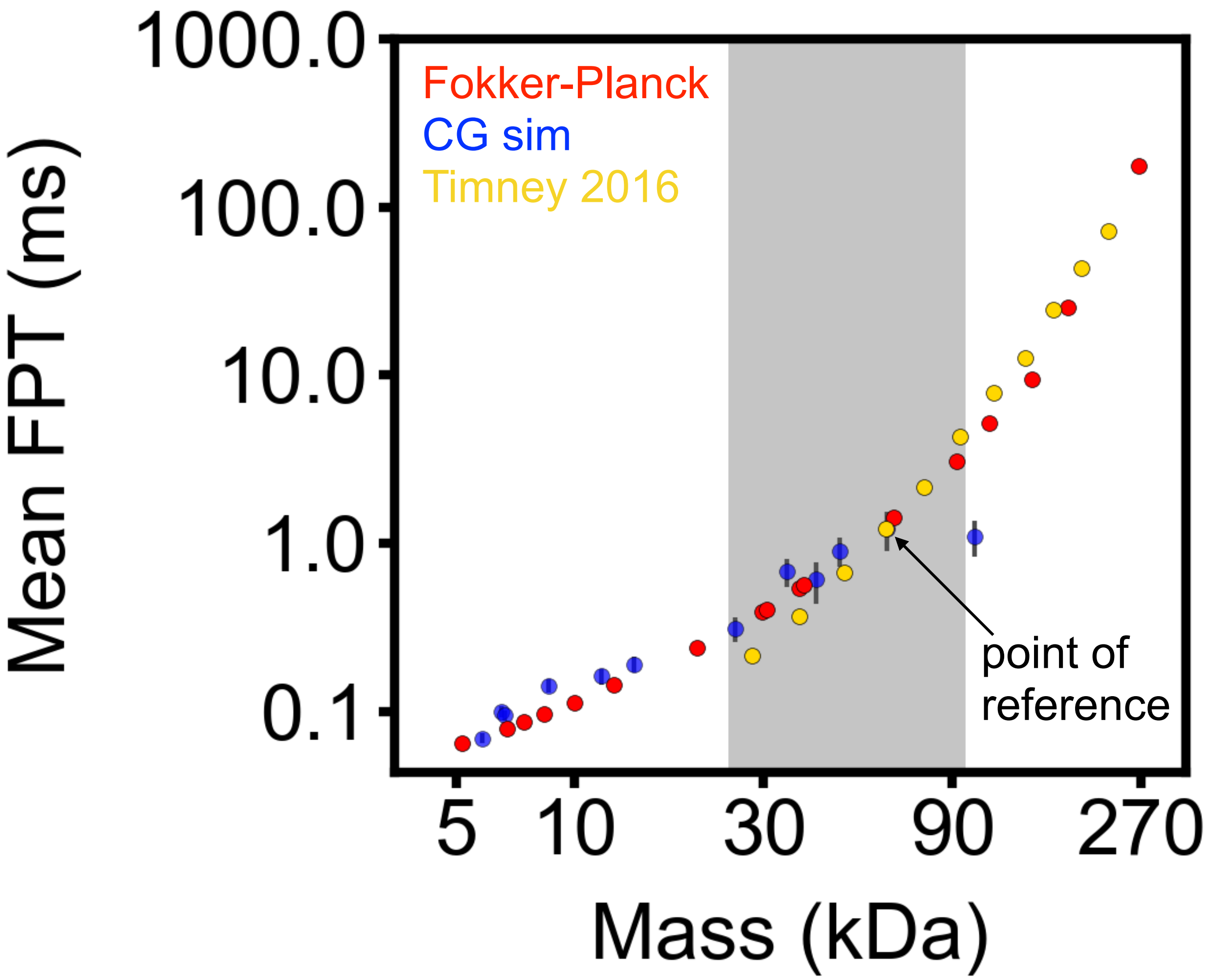


**Supplementary Fig. 12: Comparison to results of a previous simulations study.** The mean first-passage time data obtained using our Fokker-Planck approach (red) and our coarse-grained simulations (blue) are plotted as a function of molecular mass along with the simulation data reported by Timney et al., 2016 (gold). To enable quantitative comparison, the MFPT values from Timney et al. (*J. Cell Biol.*, **2016**, *215*, 57–76) were multiplied by a single constant value: the ratio of the hemoglobin's (61.5 kDa) MFPT obtained from our CG simulations and the MFPT reported by Timney et al. for a sphere of a 61 kDa mass. Note the logarithmic scale of both axes. The differences between the results of the CG simulation for smaller proteins (28 and 37 kDa) may be explained, in part, by the differences in the simulation setups, including the shape of the NPC scaffold and the FG-nup composition.

**Supplementary Table 1: Rigid body approximation of protein size and diffusion.** Starting from their crystal structures, all-atom models of the proteins were constructed to include all hydrogens atoms and used to calculated the total mass. To determine the radius of a protein, we first computed the protein's moments of inertial, $I_X$, $I_Y$, and $I_Z$. We then equated the moments to those of a constant-density ellipse, $I_X = \frac{1}{5}m(b^2 + c^2)$, $I_Y = \frac{1}{5}m(a^2 + c^2)$ and $I_Z = \frac{1}{5}m(a^2 + b^2)$, where $m$ is the total mass of the protein, and determine the length of the three semi axes $a$, $b$ and $c$. The protein radius was determined by equating the volume of a sphere to the volume of the ellipse, i.e., as $(abc)^{1/3}$. The standard deviation of the protein radius reported in the table is the standard deviation of $a$, $b$ and $c$ from that radius. The average diffusion coefficient was calculated by averaging the diagonal elements of the diffusion matrix returned for each protein by the HYDROPRO server.

| Name | PDB ID | Molecular mass (kDa) | Protein radius (Å) | Average diffusion coefficient (Å$^2$/ns) |
|---|---|---|---|---|
| Insulin | 2HIU | 5.812 | 12.61 ± 2.32 | 15.218 |
| Aprotinin | 4PTI | 6.524 | 12.88 ± 4.02 | 14.816 |
| Z-domain | 2SPZ | 6.638 | 13.07 ± 5.07 | 14.279 |
| Ubiquitin | 1UBQ | 8.565 | 14.72 ± 2.70 | 13.721 |
| Thioredoxin | 1F6M | 11.654 | 15.94 ± 2.38 | 12.456 |
| $\alpha$-lactalbumin | 1F6S | 14.066 | 17.08 ± 4.61 | 11.540 |
| GFP | 1EMA | 25.379 | 21.13 ± 4.60 | 9.462 |
| PBP | 2ABH | 34.421 | 23.38 ± 8.34 | 8.339 |
| MBP | 1ANF | 40.570 | 25.18 ± 6.70 | 7.949 |
| Enolase | 6ENL | 46.623 | 26.61 ± 5.70 | 7.640 |
| Hemoglobin | 4HHB | 61.477 | 29.85 ± 4.16 | 6.933 |
| RAR | 1G5Y | 102.044 | 35.94 ± 9.04 | 5.689 |
| GAPDH | 1U8F | 143.471 | 40.94 ± 1.62 | 5.166 |

**Supplementary Table 2: The radius and the diffusion constants of each spherical probe used for void analysis and Fokker-Planck calculations.** To enable direct comparison of the CG simulation results with the void analysis data, each spherical probe was assigned an equivalent molecular mass using the best fit power-law dependence as shown in Supplementary Fig. 7.

| Void probe radius (Å) | Diffusion coefficient (Å$^2$/ns) | Equivalent molecular mass (kDa) |
|---|---|---|
| 11.043 | 17.192 | 5.183 |
| 12.253 | 15.618 | 6.729 |
| 12.75 | 15.056 | 7.436 |
| 13.352 | 14.428 | 8.349 |
| 14.314 | 13.530 | 9.943 |
| 15.708 | 12.416 | 12.557 |
| 19.051 | 10.390 | 20.386 |
| 22.162 | 9.036 | 29.806 |
| 22.395 | 8.948 | 30.600 |
| 24.1425 | 8.348 | 36.956 |
| 24.384 | 8.272 | 37.890 |
| 30.044 | 6.822 | 64.000 |
| 34.745 | 5.964 | 92.200 |
| 37.5 | 5.558 | 111.675 |
| 41.318 | 5.082 | 142.463 |
| 45.0 | 4.698 | 176.525 |
| 52.94 | 4.042 | 265.482 |

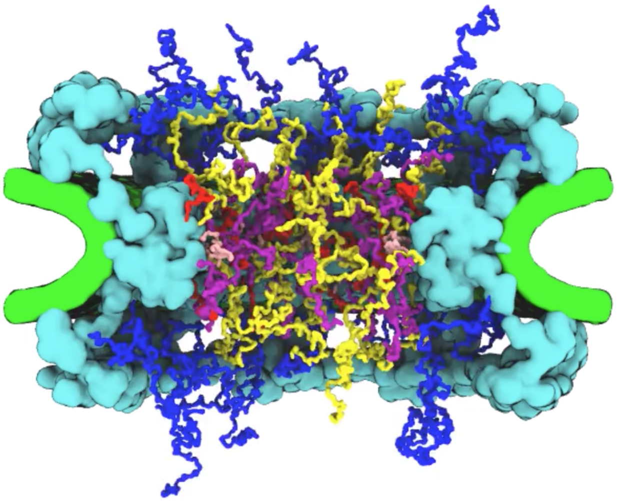

**Supplementary Movie 1: Equilibration simulation of the nuclear pore complex.** The animation illustrates the last 1,000 microseconds of a 7,500-microsecond CG simulation. Colors identify the NPC scaffold (cyan), envelope (green), and tethered FG-nups (many colors). FG-nups are colored by species.

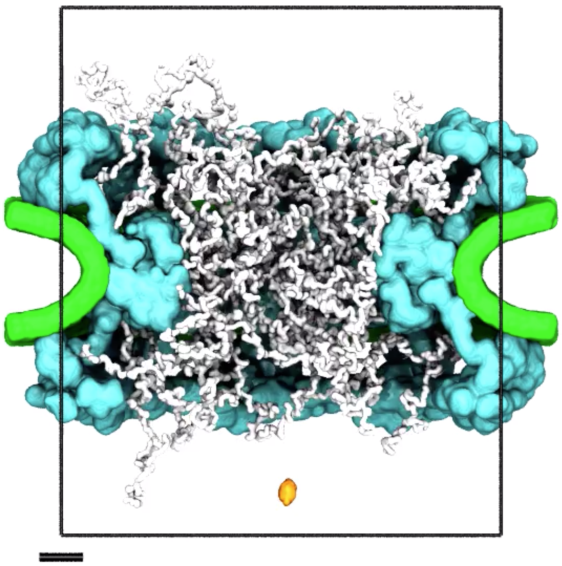

**Supplementary Movie 2: Coarse-grained simulation of passive diffusion**. The animation illustrates a 750-microsecond fragment of a CG simulation where $\alpha$-lactalbumin attempts to and eventually crosses through the NPC. The crossing even happens toward the end of the animation. Colors identify the NPC scaffold (cyan), envelope (green), tethered FG-nups (white) and $\alpha$-lactalbumin (orange). The cylindrical confinement potential, radius of 50 nm and height of 120 nm, is outlined in black. Lower-left black scale bar, 10 nm.

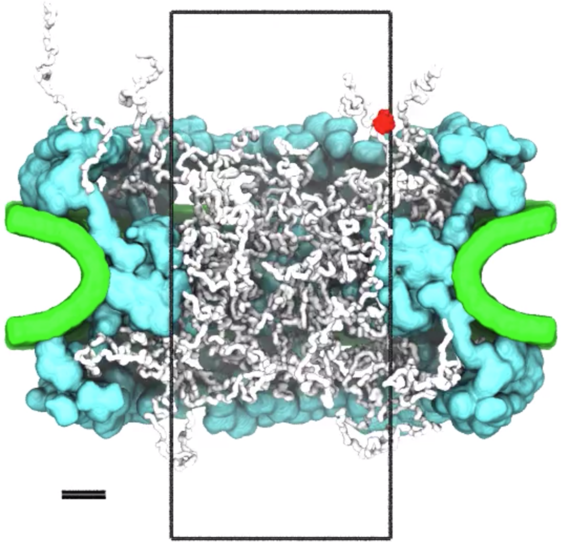

**Supplementary Movie 3: Illustration of a typical crossing event.** The animation illustrates a 6-microsecond fragment of a much longer MD trajectory where a maltose-binding protein is seen to pass through the NPC. Colors identify the NPC scaffold (cyan), envelope (green), tethered FG-nups (white) and maltose-binding protein (red). The instantaneous coordinates of the FG-nups and of the protein were averaged over two consecutive frames (20 ns) to smooth the representation of their motion. The cylindrical confinement potential, radius of 25 nm and height of 120 nm, is outlined in black. Lower-left black scale bar, 10 nm.
